# Formation of flavone-based wooly fibres by glandular trichomes of *Dionysia tapetodes*

**DOI:** 10.1101/2020.10.06.320911

**Authors:** Matthieu Bourdon, Josephine Gaynord, Karin Müller, Gareth Evans, Simon Wallis, Paul Aston, David R. Spring, Raymond Wightman

## Abstract

*Dionysia tapetodes*, a small cushion-forming mountainous evergreen in the Primulaceae, possesses a vast surface-covering of long silky fibres forming the characteristic “wooly” farina. This contrasts with some related *Primula* which instead possess a powdery farina. Using a combination of cell biology and analytical chemical techniques, we provide a detailed insight of wooly farina formation by glandular trichomes that produce a mixture of flavone and substituted flavone derivatives, including hydroxyflavones. Conversely, our analysis show that the powdery form consist almost entirely of flavone. The wooly farina in *D. tapetodes* is extruded through specific sites at the surface of the glandular head cell, characterised by a small complete gap in the plasma membrane, cell wall and cuticle. The data is consistent with formation and thread elongation occurring from within the cell. The putative mechanism of wool thread formation and its stability is discussed.

## Introduction

The genus *Dionysia* contains 55 species, found across central Asia. They are very closely related to *Primula* and some species have historically moved backwards and forwards between the two genera (Lidén 2007). Under the Angiosperm Phylogeny Project, *Dionysia* has been subsumed into *Primula* (https://www.mobot.org). However, the current Missouri/Kew “Plant list”, a working list of all plant species, still recognises *Dionysia* as a genus in its own right (http://www.theplantlist.org). Species of *Dionysia* are dwarf shrubs or woody perennials forming loose to densely compact cushions with terminal inflorescences. They grow at high altitude in mountain regions usually on limestone cliffs although granite, sandstone and dolomitic sites have also been recorded and are found in shaded or semi shaded conditions. *Dionysia tapetodes* (Bunge 1871) is the most widely distributed *Dionysia* with a range from the Kopet Dagh through NE Iran (mountains of Khorasan Province) to the mountains of Afghanistan (Grey-Wilson 1989). This wide range helps to explain the variation within the species. They form large, rather flat cushions with yellow flowers in the wild and in cultivation (Beckett *et al*. 1993).

For some species of *Primula* and *Dionysia*, a mealy deposit termed farina (latin “meal” or “flour”) is found to cover all or a subset of aerial parts of the plant. It is readily observed, for example on the leaf surface, either as a powder or, in some *Dionysia* species, as fine thread (or wool)-like fibres that are commonly referred to as “wooly farina”. Like *Primula*, in *Dionysia* both farinose and efarinose forms can exist within the same species. The powdery farina of *Primula* species is mostly comprised of 2-phenyl-4H-chromen-4-one, more commonly known as flavone (Müller 1915; Blasdale 1945). Chemical analysis of solvent-rinsed plant organs reveal a growing number of additional substituted flavones, particularly hydroxy- and methoxy-derivatives. The number of different flavone types can be extensive and their presence/absence can vary between closely-related species (Valant-Vetschera *et al*. 2009, 2010; Bhutia *et al*. 2013; Hinterdobler *et al*. 2017). Given these flavone assignments represent total surface-extracted flavones that include different tissue/cell types, it is not clear which substituted flavones actually constitute the farina of a given species.

Farina biosynthesis takes place in glandular trichomes that typically consist of a stalk attached to a single round cell known as the glandular head (Fico *et al*. 2007; Vitalini *et al*. 2011; Bhutia and Valant-Vetschera 2012). The crystals of powdery farina are seen to coat the head cell and, at the subcellular level, a proliferation of smooth ER is observed suggesting a site of synthesis and transport of some biosynthetic intermediates (Wollenweber and Schnepf 1970; Gunning and Steer 1996). Part of the biosynthetic pathway to flavones is known but for synthesis of more complex derivatives is less well understood (Jiang *et al*. 2016). Chalcone synthase (CHS) represents the first step in the flavonoid biosynthetic pathway and immunolocalization of the enzyme in farinose *Primula kewensis* shows high signal in the gland head cell (but not in an *efarinose* mutant), suggesting the full flavone farina biosynthetic machinery is present in this cell (Schopker *et al*. 1995). Furthermore, immunogold labelling showed enrichment of CHS in spherical bodies in the cytoplasm. As flavone farina biosynthesis continues, the subcellular locations of intermediates and final products are not known. Generally, flavone glycones, being soluble, can accumulate in the vacuole (Marinova *et al*. 2007) but, for farina producing plants, the storage location of the insoluble aglycone is not well understood, possibly being localised to a modified vacuole-type organelle that can stretch across the cell (van Brederode and Steyns 1985). These flavone and flavone-type aglycones need to be transported out of the cell and through the cell wall and cuticle for deposition on the surface of the hair cell. Presumably this becomes more complicated for wooly farina whereby the flavone building blocks need to be concentrated to a single exit site to produce an elongating fibre. We present here our results showing wooly farina composition and the subcellular organisation of the glandular trichome head cell that is the site of farina synthesis in *D. tapetodes*. We identify the wool as a mixture that includes flavone and hydroxyflavones that emerge from the head cell and is threaded through distinct gaps in the cell wall and cuticle. The mechanism of wool formation is discussed.

## Materials and Methods

### Plant propagation and harvesting

We used *Dionysia tapetodes* accession 20140435 that was received from the Royal Botanic Garden Edinburgh (accession 19822508) comprising wild material collected by Prof. T.F. Hewer (number 1164) between 1969 and 1971. *D. tapetodes* was grown at Cambridge University Botanic Garden (Cambridge, UK) in clay pots plunged in sand in an alpine house to keep the roots cooler and at a more stable temperature. The potting compost comprised 50% loam-based compost, 30% 1-9mm grit, 10% sharp sand, 10% seramis and a small amount of slow release fertiliser. The compost is top-dressed with grit which is carefully worked under the collar of the plant to reduce the risks of basal rot. Given that *Dionysia* are prone to rosette burning from overexposure to sunlight, a temporary shading screen was placed over the collection from mid-March and removed later in the season after sun intensity decreased. Old flowers were carefully removed to prevent any botrytis infection moving from dead flowers into living material. Watering adhered to a strict regime: Young plants were watered from underneath via a water bath and once they passed one year old were then watered overhead without getting the cushion surface wet. Water was applied sparingly in the winter months and increased once the plants were in active growth.

### Cryo-Scanning Electron Microscopy (cryoSEM) and cryo-fracture

Rosettes of 5-8 leaves were mounted, frozen in nitrogen slush, platinum coated and fractured as previously described (Wightman *et al*. 2018). To accommodate better fractures without dislodging trichomes, some samples were dipped in 70% v/v ethanol to remove the wool and then air-dried prior to freezing. Cryo prepared samples were viewed using a Zeiss EVO HD SEM fitted with a backscattered electron detector and 25 kV acceleration voltage.

### Low kV SEM imaging of wool fibres

Clumps of uncoated wool fibres were placed on a sticky carbon tab and mounted on an SEM stub in the Zeiss EVO HD SEM at high vacuum and 1kV accelerating voltage using SE detector fast scanning with frame averaging to prevent wool movement.

### Embedding, sectioning and imaging of trichomes by transmission electron microscopy (TEM) and light (epifluorescence) microscopy

Leaves were dissected with razor blades in a solution consisting of 4% formaldehyde (freshly prepared from paraformaldehyde powder, Sigma) and 0.5% glutaraldehyde (Sigma) in PBS buffer. Fixation, dehydration, resin infiltration and antibodies washing steps were all microwave (MW) assisted using a PELCO BioWave Pro (Ted Pella). Fixation was realized at 150W, under vacuum (20Hg) (5x 1min). Samples were left in the fixative overnight at 4°C and then washed 3 times in PBS. Stems were then parallelly aligned and embedded in 1% agarose in PBS. Samples were then processed through increasing dehydration steps (25%, 50%, 70%, 90%, 96%, 3x 100% ethanol, vacuum 20Hg, MW 150W 5min), and left overnight in 100% ethanol at 4°C. Resin infiltration (LR White medium grade, Agar scientific) was then realized through increasing resin concentration: 33% Resin in ethanol 100%, 66% Resin in ethanol 100%, and 3 times 100% Resin (20Hg, MW 200W 5min). Samples were left at least 24h in 100% resin for effective penetration in the samples. Resin polymerization was subsequently realized at 60°C during 18h.

For epifluorescence microscopy, semi-thin sections (1μm) of the whole leaf bud were then obtained with a Leica EM UC7 ultramicrotome using a Histo Jumbo 8mm diamond knife (DiATOME) and laid on droplets of sterile water on uncoated glass microscopy slides. In order to avoid folds on sections during the water drying process, slides were dried on a hot plate at 55°C (Leica). Slides were finally mounted in a 1:1 solution of AF1 antifadent (Citifluor) with PBS, containing calcofluor as a cell wall counterstaining, and imaged by confocal laser scanning microscopy (Zeiss LSM700).

For TEM observations, 100nm thin sections were obtained with a Leica EM UC7 ultramicrotome using an ultra 45° diamond knife (DiATOME) and deposited on 200-300μm mesh formvar-coated nickel grids

Samples were post-stained using Reynold’s lead citrate and uranyl acetate for 3 min in each. Thin sections were viewed in a FEI/Thermofisher Tecnai G20 electron microscope run at 200 keV with a 20 μm objective aperture to improve contrast. Images were taken with an AMT camera running DEBEN software.

### Field Emission Scanning Electron Microscopy (FE-SEM) of leaf sections

Whole rosettes were dissected with razor blades to remove apical parts of leaves and immediately submerged in fixative (2 % glutaraldehyde/2 % formaldehyde in 0.05 M sodium cacodylate buffer pH 7.4 containing 2 mM calcium chloride) under vacuum overnight at room temperature. After washing 5x in DIW (deionised water), samples were osmicated (1 % osmium tetroxide, 1.5 % potassium ferricyanide in 0.05 M sodium cacodylate buffer pH 7.4) for 3 days at 4°C. Then samples were washed 5x with DIW and treated with 0.1 % (w/v) thiocarbohydrazide/DIW for 20 minutes at room temperature in the dark. After washing 5x in DIW, samples were osmicated a second time for 1 hour at RT (2% osmium tetroxide/DIW). After washing 5x in DIW, samples were blockstained with uranyl acetate (2 % uranyl acetate in 0.05 M maleate buffer pH 5.5) for 3 days at 4°C. Samples were washed 5x in DIW and then dehydrated in a graded series of ethanol (50%/70%/95%/100%/100% dry) 100% dry acetone and 100% dry acetonitrile, 3x in each for at least 5 min. Samples were infiltrated with a 50/50 mixture of 100% dry acetonitrile/Quetol resin (without BDMA) overnight, followed by 3 days in 100% Quetol (without BDMA). Then, the sample was infiltrated for 5 days in 100% Quetol resin with BDMA, exchanging the resin each day. The Quetol resin mixture is: 12 g Quetol 651, 15.7 g NSA, 5.7 g MNA and 0.5 g BDMA (all from TAAB). Samples were placed in embedding moulds and cured at 60°C for 3 days. Semi-thin sections (1 μm) of the whole leaf bud were then obtained with a Leica EM UC7 ultramicrotome using a Histo-Jumbo 8mm diamond knife (DiATOME) and laid on droplets of sterile water on uncoated glass microscopy slides. In order to avoid folds on sections during the water drying process, slides were dried on a hot plate at 55°C (Leica). Glass slides were then resized with a glass knifemaker to be mounted on aluminium SEM stubs using conductive carbon tabs, and the edges of the slides were painted with conductive silver paint. Samples were sputter coated with 30 nm carbon using a Quorum Q150 TE carbon coater. Samples were imaged in a Verios 460 scanning electron microscope (FEI/Thermofisher) at 4 keV accelerating voltage and 0.2 nA probe current in backscatter mode using the concentric backscatter detector (CBS) in immersion mode at a working distance of 3.5-4 mm; 1536 x 1024 pixel resolution, 3 us dwell time, 4 line integrations. Stitched maps were acquired using FEI MAPS software using the default stitching profile and 10% image overlap.

### Measurements of wool fibre diameter

Images of wool fibres, attached to leaves of an isolated *D. tapetodes* rosette, were taken with a Keyence VHX-7000 microscope at 2500x magnification and illuminated with full field coaxial light. 2D depth-up mode was used for in-focus acquisitions. Fibre width measurements were carried out using the point-to-point measuring tool in the Keyence software.

### Raman microscopy of farina

Raman microscopy was carried out on a Renishaw InVia instrument equipped with a 785 nm laser. *D. tapetodes* farina wool or *P. marginata* powder was carefully placed on a quartz slide and brought in to focus under a 50x dry objective lens. Raman acquisitions used a 1200 l/mm grating, 1200 cm^-1^ centre, 785 nm laser at 10% power, regular confocal mode and 4 s exposure with 3 accumulations. At least 3 spectra per sample were averaged in order to improve signal-to-noise. To find close matches with reference Raman spectra, the experimental spectra were used as a search input against the Raman databases, that include some flavone derivatives, in the KnowItAll software (Bio-Rad Inc.) using the “SearchIT” tool and then candidate spectra were visualised by eye to remove false positives. Both default and Euclidean distance search settings were used. Matches are ranked according to their hit quality index (out of a maximum of 100). *P. marginata* farina gave a close match (97/100) with flavone. *D tapetodes* gave no close matches but yielded good correlation (70-80/100) to reference spectra of hydroxy- and methoxy-flavone derivatives that included 7, 2’-dimethoxyflavone; 3,7-dimethoxyflavone and 6-, 7-, or 8-hydroxy-derivatives. For fastFLIM-Raman correlative imaging of leaves submerged in water (Supplementary data Fig. S9) a confocal-Raman microscope, described in (Wightman *et al*. 2019), used the following settings: 25x 0.95 NA water dipping objective lens, FLIM 440 nm pulsed laser (at 20 MHz) with detector window set between 448 nm and 511 nm and 80 iterations. Raman: 1200 l/mm grating, 1200 cm^-1^ centre, 785 nm laser, 50% power, 15 s exposure with 2 accumulations used in line scan mode that intersected a glandular head cell.

### Reagents, solvents and sample preparation for chemical analysis

Pure Flavone was purchased from Alfa Aesar as a white solid with 99% purity (CAS no. 525-82-6, catalogue no. A13627) and used without further purification. All solvents were anhydrous and used as purchased without any further purification. Flavone wool from was picked from *D. tapetodes* leaf surfaces using fine tweezers and placed in a microcentrifuge tube. The wool sample (approximately 0.5 mg) was dissolved in 50 μL of acetonitrile/water (1:1) with a few drops of dimethylsulfoxide (DMSO) to aid solubility. Sample preparation was performed in this way for analytical HPLC, LCMS and HRMS analysis.

### Analytical high-performance liquid chromatography (HPLC)

Analytical HPLC was run on an Agilent 1260 Infinity using a Supelcosil ABZ+PLUS column (150 mm × 4.6 mm, 3 μm) eluting with a linear gradient system (solvent A: 0.05% (v/v) trifluoroacetic acid (TFA) in H2O, solvent B: 0.05% (v/v) TFA in acetonitrile (MeCN)) over 15 min at a flow rate of 1 mL/min.

### Liquid chromatography mass spectrometry (LCMS)

Chromatographs were recorded on a Waters ACQUITY H-Class UPLC with an ESCi Multi-Mode ionisation Waters SQ Detector 2 spectrometer (LC system: solvent A: 2 mM ammonium acetate in water/MeCN (95:5); solvent B: 100% MeCN; column: AQUITY UPLC CSH C18, 2.1*50 mm, 1.7 μm, 130 Å; gradient: 63 5-95% B over 3 min with constant 0.1% formic acid).

### High resolution mass spectrometry (HRMS)

HRMS was carried out on a Waters LCT Premier Time of Flight mass spectrometer. ESI refers to the electrospray ionisation technique.

### Nuclear magnetic resonance (NMR) spectroscopy

All pulse sequences are the default (with the exception of the DEPT135) from the Topspin 3.2pl7 software used to control the acquisition. The analysis required ^1^H, ^13^C, DEPT135, DFQ-COSY, Heteronuclear Single Quantum Coherence (HSQC, with DEPT 135 editing) and Heteronuclear Multiple Bond Correlation Spectroscopy (HMBC) spectra. All necessary shaped and decoupling pulses were calculated by the software, using defined 90 degree pulses.

^1^H NMR: Proton magnetic resonance spectra were recorded using an internal deuterium lock (at 298 K unless stated otherwise) on Bruker DPX (400 MHz; ^1^H-^13^C DUL probe), Bruker Avance III HD (400 MHz; Smart probe), Bruker Avance III HD (500 MHz; Smart probe) and Bruker Avance III HD 62 (500 MHz; DCH Cryoprobe) spectrometers. Pulse sequence used zg30 – PLW1=14W, P1=10.5μs, SW=20 ppm, TD=64 K, AQ=3.28 s, D1=1 s, NS=16. Proton assignments are supported by ^1^H-^1^H COSY, ^1^H-^13^C HSQC or ^1^H-^13^C HMBC spectra, or by analogy. Chemical shifts (δH) are quoted in ppm to the nearest 0.01 ppm and are referenced to the residual non-deuterated solvent peak. Discernible coupling constants for mutually coupled protons are reported as measured values in Hertz, rounded to the nearest 0.1 Hz. Data are reported as: chemical shift, multiplicity (br, broad; s, singlet; d, doublet; t, triplet; q, quartet; m, multiplet; or a combination thereof), number of nuclei, coupling constants and assignment.

^13^C NMR: Carbon magnetic resonance spectra were recorded using an internal deuterium lock (at 298 K unless stated otherwise) on Bruker DPX (101 MHz), Bruker Avance III HD (101 MHz) and Bruker Avance III HD (126 MHz) spectrometers with broadband proton decoupling. 1024 scans (NS) were acquired using pulse sequence ‘zgpg30’, with waltz16 1H decoupling. 90 degree 13C pulse set to 21W (PLW1) for 9.5 μs (P1). 209786 points (TD) were digitised over 3.02 s (AQ), relaxation delay set to 2 s (D1). Sweep width was 276 ppm (SW), with an irradiation frequency of 110 PPM (O1P). Carbon spectra assignments are supported by DEPT editing, ^1^H-^13^C HSQC or ^1^H-^13^C HMBC spectra, or by analogy. Chemical shifts (δC) are quoted in ppm to the nearest 0.1 ppm and are referenced to the deuterated solvent peak. Data are reported as: chemical shift, number of nuclei, multiplicity, coupling constants and assignment. Magnetic resonance spectra were processed using TopSpin (Bruker). An aryl, quaternary, or two or more possible assignments were given when signals could not be distinguished by any means. Standard flavone numbering was followed.

DEPT 135: Pulse sequence dept135sp – This is a minor modification of the DEPT sequence to optimise the spectral baseline and uses an adiabatic shape for 180 degree carbon pulses. Carbon pulse powers as ^13^C experiment above, SW=236.7 ppm, TD=65536, AQ=1.10 s, D1=2 s, O1=100 ppm, NS=64. Waltz16 decoupling.

DFQCOSY: This is a double-quantum filtered experiment; using gradient selection; pulse sequence cosygpmfqf. Non-uniform sampling; using a Poisson-gap weighted schedule was used to acquire 37.5% of 512 increments, each with 2 scans (SWF2=13.37 ppm, TD=4k, AQ=0.31 s, D1=2 s). Proton pulse powers as above. Processed to 2k x2k points using a sine function (SSB=2.5)

HMBC: This experiment is phase sensitive; uses Echo/Antiecho gradient selection, with a three-fold low-pass J-filter to suppress one-bond correlations; pulse program ‘hmbcetgpl3nd’. Acquired in phase sensitive mode using Echo/Antiecho-TPPI gradient selection, with a 3 step low pass j-filter to suppress 1 bond correlations. Long-range J-JCH parameters set to 10 Hz. Non-uniform sampling; with a Poisson--gap weighted schedule was used to acquire 37.5% of 768 increments; each with 2 scans (SWF1=250, SWF2=12.02 ppm, TD=4096, AQ=0.34 s, D1=2 s). Processed to 2048×2048 using a sine function (SSB=4 & 2 for F2 and F1), then converted to magnitude mode in F2. (Topspin command ‘xf2m’)

HSQC: The HSQC was acquired using a Bruker Avance III HD 500Mhz equipped with a dual 13C/1H cryoprobe; using Topspin 3.2pl7. It was acquired in ‘non-uniform sampling’ mode and samples 25% of 1024 increments, using a ‘poisson-gap’ schedule. The data was processed using the default compressed sensing (CS) method in Topspin 3.5pl7 (on your computer) to 2048×2048 data points. This experiment is configured to give −CH2 groups an opposite phase to the −CH and −CH3 groups, using the hsqcedetgpsp.3 pulse sequence. Acquired in phase sensitive mode using Echo/Antiecho-TPPI gradient selection, multiplicity edited during selection step with shaped adiabatic pulses. Non-uniform sampling; with a Poisson-gap weighted schedule was used to acquire 25% of 1024 increments; each with 2 scans (SWF1=190 ppm, SWF2=12.99 ppm, TD=1816, AQ=0.14 s, D1=0.8 s). Carbon and proton pulse powers as above. Processed to 2048×2028 points using a qsine function (SSB=2). Standard flavone numbering:

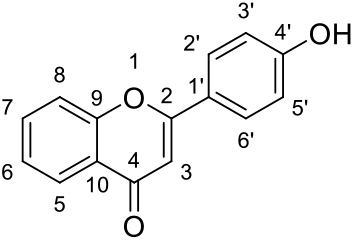

## Results

### Wooly farina fibres of *Dionysia tapetodes* have distinct surface grooves and differ in composition to powdery farina

*D. tapetodes* grows as a densely packed cushion (Fig. 1A) with large quantities of “wooly” fibres observed on both the adaxial and abaxial surface of the *D. tapetodes* leaf (Fig. 1B). Wooly farina production likely occurs throughout leaf growth since it is present in both young leaves (found at the centre of the rosette) and older leaves (positioned on the outside of the rosette). Farina production at early stages of growth are consistent with the observed entanglement of fibres between neighbouring leaves as revealed by scanning electron microscopy (SEM) (Fig. 1C). Individual wooly fibres range in width from 0.9 to 2.1 microns, with a mean width of 1.6 microns (n=51). High magnification low kV SEM of uncoated fibres shows fine grooves on their surface, making a series of ridges arranged longitudinally (Fig. 1D). In order to understand how these fibres are formed and maintained, we compared the Raman spectra of wooly farina of *D. tapetodes* with the powdery farina found near the leaf margin of *Primula marginata*. Figure 1E shows an overlay of the spectra from both farina types. The powder spectrum of *P. marginata* matches that of pure flavone (see materials and methods for details of spectra correlation), indicating that flavone makes up all or the vast majority of the farina in this species. The spectrum of *D. tapetodes* wooly farina, however, only shares some of the peaks with the flavone powder of *P. marginata;* at 674, 1001, 1012 and 1568 cm^-1^. This suggests a substituted and/or mixture of flavones comprise the wooly farina of *D. tapetodes*. Spectral search and correlation software were used to identify candidate functional groups (see materials and methods) based on the acquired Raman spectra and compared to a reference database that included some flavone derivatives. While *P. marginata* farina correlated very highly with unsubstituted flavone (hit quality index 97 out of a possible 100 maximum), *D. tapetodes* wooly farina gave no close matches. However, the best hits (hit quality index 70-80) were to various combinations of mono-and di-hydroxy- and methoxy-substituted flavones. This suggests that hydroxy- and/or methoxy functional groups might co-exist with flavone in the wool fibres.

**Figure 1.**
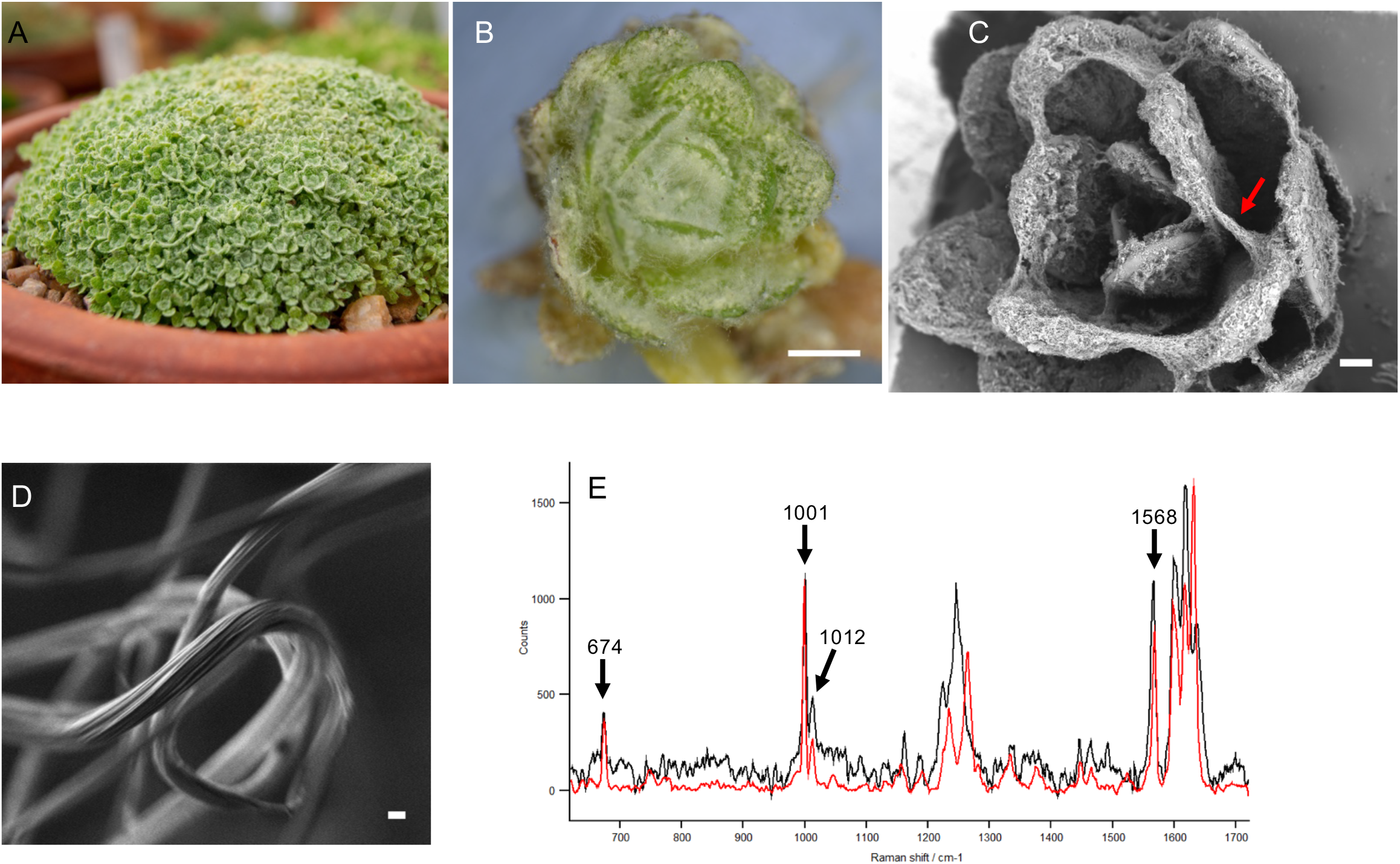
Farina wool observations on leaves of *Dionysia tapetodes*. A) Overview of the densely-packed, cushion-forming *D. tapetodes*. B) Stereomicroscope image of white wooly farina on leaves of *D. tapetodes*. Scale bar = 1 mm. C) Scanning electron microscopy (SEM) of wooly farina distributed on and between leaves (e.g. red arrow) of *D. tapetodes*. Scale bar = 300 μm. D) High magnification SEM image of farina fibre showing fine surface structure. Scale bar = 1 μm. E) Raman microscopy of fingerprint region of farina of *Dionysia tapetodes* (black) compared with that of the powdery farina from *Primula marginata* (red). Principle peak assignments (cm^-1^) common to both types of farina are indicated.

### The wooly farina of *D. tapetodes* comprises mostly flavone mixed with substituted flavones

A sample of the wooly farina from *D. tapetodes* was further analysed by high-performance liquid chromatography (HPLC), liquid chromatography-mass spectrometry (LCMS), high-performance mass spectrometry (HRMS) and nuclear magnetic resonance (NMR) spectroscopy. All data are presented together with a detailed interpretation in Supplementary dataset S1 that additionally comprises Supplementary data Figs. S1–S7 and Table S1. This analysis revealed that the wooly farina sample was mainly composed of unsubstituted flavone (>90% by analytical HPLC, structure A in Fig. 2). Identical analytical data was obtained for a pure sample of flavone from a commercial source (Supplementary data dataset S1, Fig. S4). The HPLC, LCMS and HRMS data indicated that there are other, substituted flavone species present in the wooly farina sample; exactly how many other species is not clear. HPLC analysis gave only one, whereas LCMS data suggested the presence of multiple other flavonoids. Mass ions were observed in the LCMS corresponding to hydroxy- and methoxy-substituted flavones (Supplementary data Figs. S1 and S2). To provide more information on the structure of the minor flavone species, 2D NMR was carried out and yielded candidate structures corresponding to 2’-hydroxyflavone and 4’-hydroxyflavone (Fig. 2, Supplementary data Fig. S7).

**Figure 2.**
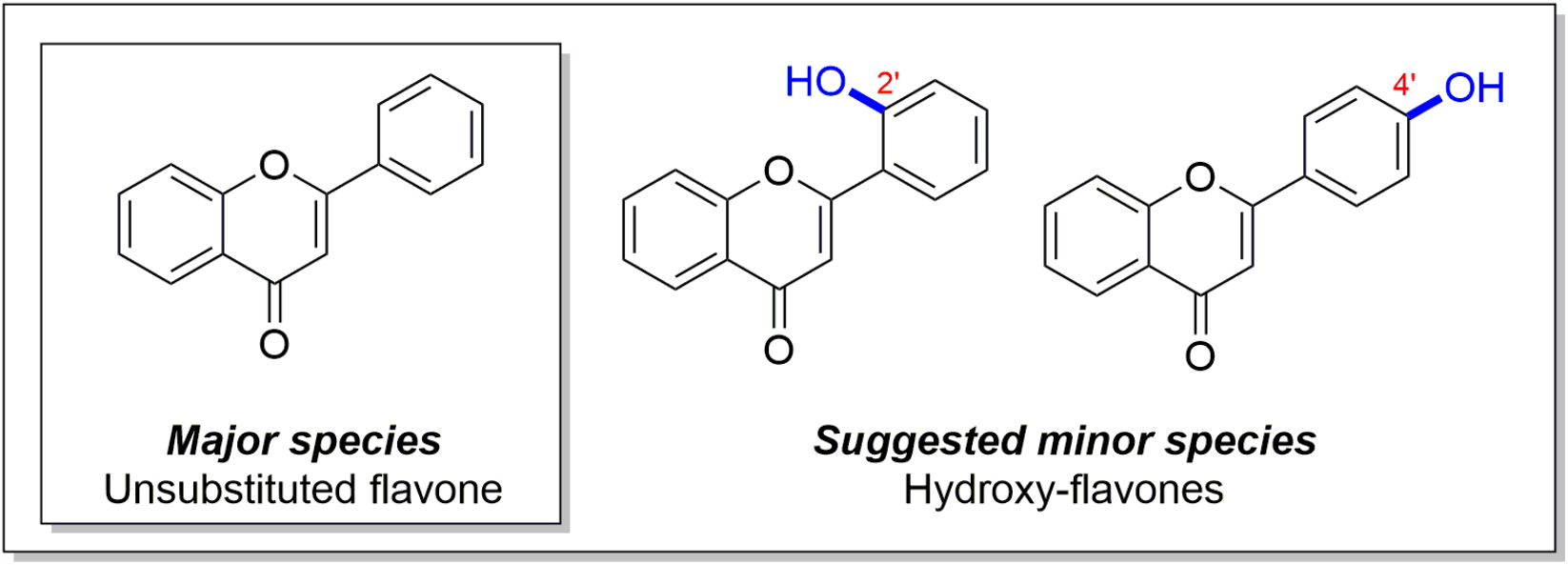
Molecular structures of candidate molecular species that form the wooly farina of *D. tapetodes*. Structures are derived from chemical analyses described in dataset S1.

### The wool fibres emerge from distinct holes in the cell wall of the glandular trichome head cell and are in close proximity to dense material of the vacuole

Focal points representing the origin of numerous fibres can be seen on the *D. tapetodes* leaf surface (Fig. 3A) corresponding to mature flattened glandular trichomes (Fig. 3B). Interspersed between these locations are numerous smaller and extended trichomes that have fewer, single or no wool fibres emerging from the head cells and likely represent glandular trichomes at different levels of maturity (Fig. 3B and 3C). It is evident that, when looking at the smaller trichomes, farina emerges from a few discrete locations on the glandular head cell that may grow in number as the cell ages. This appears to be in contrast to *P. marginata*, where smaller crystals completely cover the circumference of the head cell (Fig. 3D). These data show that, for *D. tapetodes*, flavone incorporation and extrusion needs to be sustained at the same site on the (head) cell surface. We used cryoSEM to observe, close-up, the junction between wool fibre and cell (Fig. 4A, B) where the fibre appears to come out of the cell wall. We generated random cryofractures through the leaf that might include a glandular head cell, however, the interaction of the fracture knives with the wool resulted in dislodging of the trichome from the leaf, as well as more cellular debris at the fracture site. We found a 5 s wash with 70% v/v ethanol removed the surface wool and resulted in better cryofractures with less debris. Sites of wool fibre extrusion, represented as small “craters” on the cell surface, are visible after the washing (Fig. 4C). Cryofractures that included any part of the gland head cell were, however, very rare. One fracture removed most of the head cell leaving a part of the head and some cytoplasm intact (Fig. 4D).

**Figure 3.**
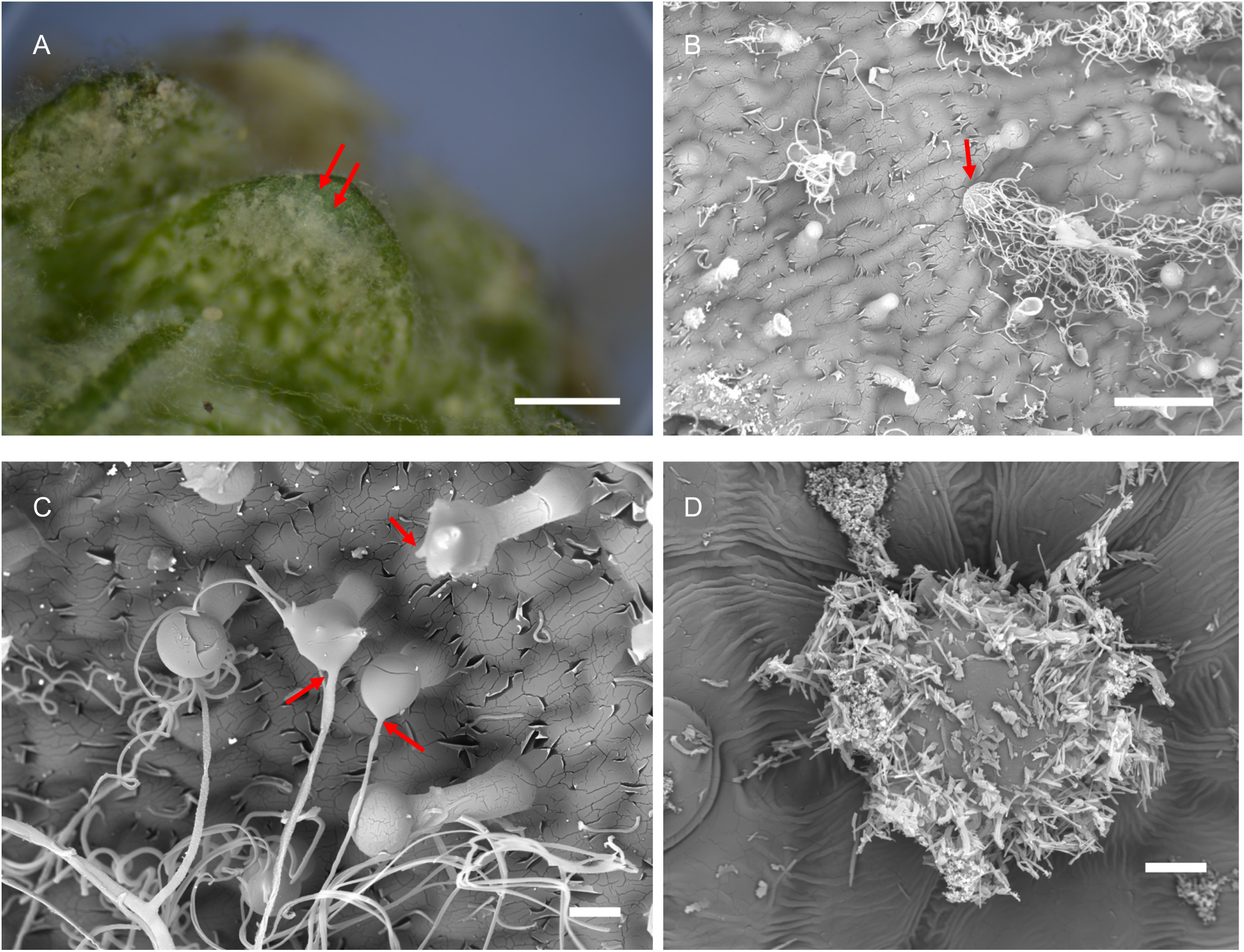
Formation of farina at the single cell level. A) Stereomicroscope image showing examples of wool exit points (red arrows) on the surface of the leaf of *D. tapetodes*. Scale bar = 500 μm. B) SEM image of glandular trichomes, including an example of a mature glandular trichome producing large quantities of wooly farina (red arrow). Scale bar = 50 μm. C) SEM image showing farina wool exit points (red arrows) from glandular trichomes. Scale bar = 10 μm. D) SEM image showing glandular trichome of *P. marginata* covered with powdery flavone farina. Scale bar = 10 μm.

**Figure 4.**
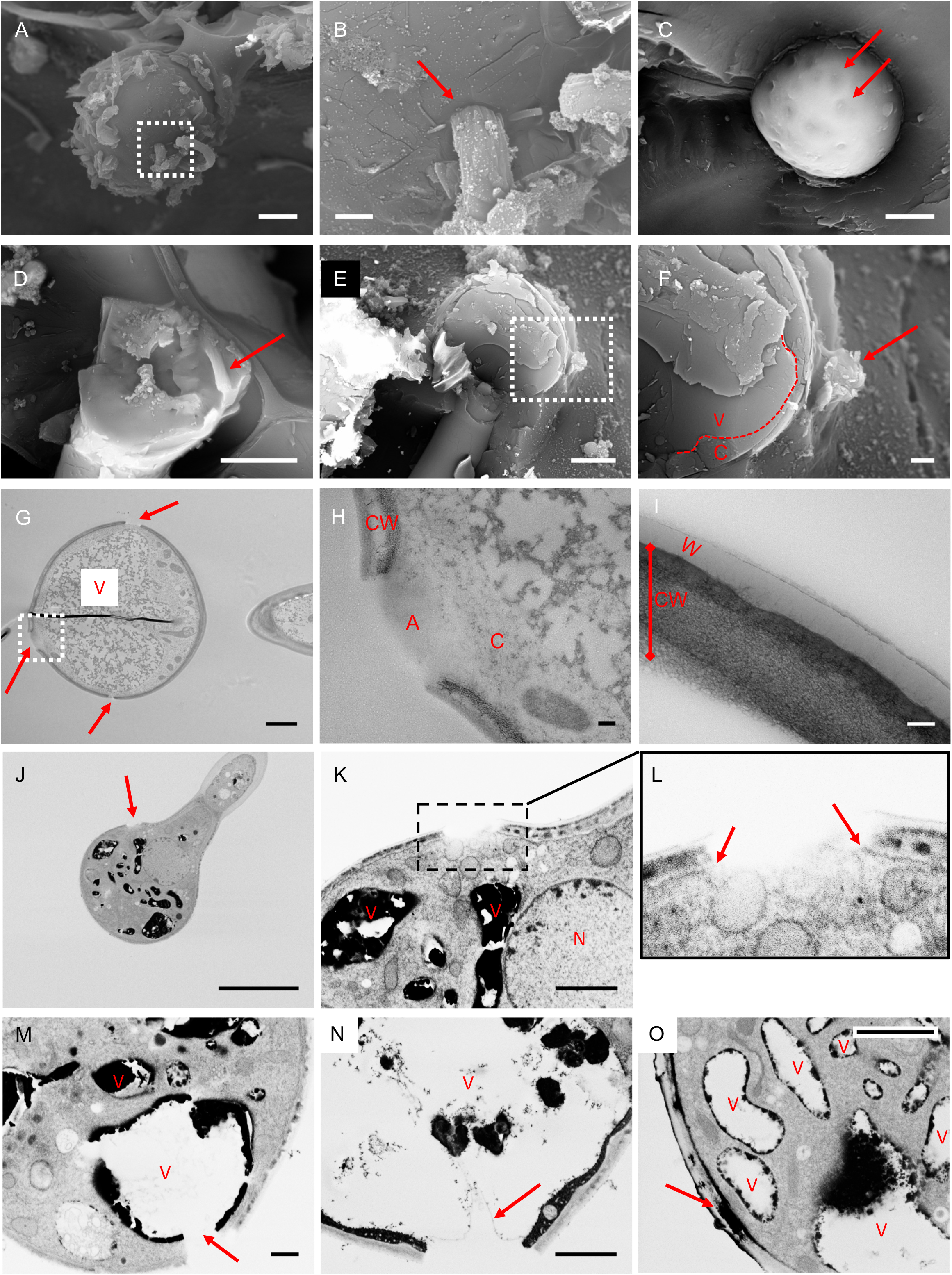
Exit points of farina wool from glandular trichomes. A) SEM image showing wool exiting from the surface of a glandular trichome cell. A magnified view of the boxed region is shown in (B). Cell surfacefarina interface is labeled by a red arrow. Bars = 5 μm (A) and 1 μm (B). C) Surface of glandular trichome cell after ethanol wash to remove farinose material. Examples of wool exit points are labeled by red arrows. Bar = 5 μm. D) SEM after cryo-fracture showing inner face of part of the surface of a trichome cell. A piece of farina wool is found to be present intact inside the cell (red arrow). Bar = 5 μm. E) SEM after cryofracture revealing cell contents. Bar = 5 μm. A magnified view of the boxed area is shown in (F) where wool exit is observed (red arrow). The dashed line represents the location of the tonoplast. V= Vacuole, C= cytoplasm. The Vacuole is seen to be in close proximity to the wool exit site. Bar = 1 μm. G) Transmission EM (TEM) image of section through glandular trichome. Wool exit sites are marked by red arrows and are defined as a gap in the cell wall. Electron dense material, consistent with vacuole contents (V), are in close proximity to these sites. Dark line in the centre is an artefact from a fold in the section. Bar = 2 μm. H) a magnified view of a wool exit site (boxed region) showing a gap in the cell wall (CW) and an amorphous region (A) within this gap. Beneath the amorphous region is observed a small amount of cytoplasm (C). Bar = 200 nm. (I) TEM image of a section of intact cell wall (CW) overlayed by a waxy low density layer (W). Bar = 100 nm. J, K, L) FE-SEM of glandular head showing wool exit site (arrow in J) Bar = 10 μm. A magnified view of the area covering the exits site is shown in (K) together with the electron dense vacuoles (V) and the Nucleus (N) Bar = 2 μm. Further magnification (L) shows the plasma membrane does not traverse the gap (arrows). M, N) Exit sites that extend to the vacuole. In (N) the tonoplast is found to be intact (arrow). Bars = 2 μm. O) A build up of electron dense material accumulates within a void in the cell wall (arrow). Vacuoles (V) containing similar electron dense material are found in close proximity. Bar = 2 μm.

Ethanol washing had removed the external wool fibres from the sample, however, a single fibre is observed inside the cell that was protected from the solvent (red arrow in Fig. 4D). This suggests the wool is made inside the cell and then somehow threaded through a gap in the wall. A second fracture was obtained through the centre of the gland hair cell (Fig. 4E) and a wool exit site could be observed close to the fracture plane. At this location the vacuole appeared to be close to the surface with a small amount of cytoplasm in between (Fig. 4F). Given that we could not obtain a fracture through the site of wool extrusion in the cell wall, we carried out Transmission Electron Microscopy (TEM) on post-stained sections of fixed glandular trichomes. TEM processing completely removes all the farina wool, however, complete gaps in the cell wall were observed (Fig. 4G) and were in close proximity to dense particulates that, based on the FE-SEM imaging described below, represent contents of the vacuole. These gaps are not an artefact from masking or non-uniform staining for TEM because they were also observed, in addition to the dense vacuoles, with light microscopy (Supplementary data Fig. S8). A small amount of cytoplasm, as well as an amorphous stained region, can be observed between the dense particulates and the cell wall gap (Fig. 4H). A magnified TEM view of the wall shows a mesh structure with layers of differing density that likely represent polysaccharide components including cellulose (Fig. 4I). A low density outer layer is present that is consistent with cuticular wax plus a thin epicuticular layer on top (Fig. 4I). This was confirmed by correlative fluorescence lifetime imaging and Raman microscopy which showed six Raman peaks (1062, 1132, 1294, 1416, 1438, 1465 cm^-1^) that correspond to the major peaks of epicuticular waxes from Sorghum (Farber *et al*. 2019). These wax peaks were specific to the outer surface of the gland hair cell and were found together with the flavone-type signals (Supplementary data Fig. S9). An intracellular store of flavone was also located.

In order to identify how the generation of a hole in the cell wall is coupled with directed deposition of flavone to make a thread, we used a higher throughput approach for imaging larger numbers of glandular trichomes. Full rosettes that include leaves at various stages of development were stained, embedded and sectioned. These large area sections were viewed by field emission scanning electron microscopy (FE-SEM) of backscattered electrons and gave good contrast for observations of membranes. Out of 11 gaps examined, where vacuoles could be easily distinguished from other organelles due to their intense staining, 6 gaps were within 1 μm of its closest vacuolar compartment with distances in the range 0 – 2.6 μm (mean 1.1 μm, median 0.72 μm). FE-SEM showed the gaps in the cell wall and cuticle also includes the plasma membrane (Fig. 4J-L,); 15/15 gaps observed with FE-SEM had no plasma membrane crossing the gap. For two glandular heads, we observed the gap to extend further inside the cell as far as the vacuole and in one section the tonoplast was intact (Fig. 4M, N). We observed fully formed trichomes where the dense staining was seen in a void within the cell wall of width 1.15 μm consistent with the size expected of a forming wool thread (Fig. 4O) and suggesting a careful coupling between wall digestion and deposition of the flavone and derivatives.

## Discussion

*D. tapetodes* is covered with wooly farina around the leaves. It is not entirely understood what the purpose of the farina is, however, tolerance to freezing and high UV have been cited (Caldwell 1971; Sisa *et al*. 2010). Consistent with the latter, there exists an efarinose *D. tapetodes* accession at the Cambridge University Botanic Garden and horticulturists working closely with the plants report that it scorches more easily in the sun compared to the farinose accessions.

The wooly farina threads radiate in all directions from mature glandular head cells of trichomes that are interspersed on the leaf epidermis. In addition to biosynthesis, these head cells appear to act as anchor points of the wool which can extend between adjacent leaves. The study of wool farina formation in *D. tapetodes* is made challenging by the physical and chemical characteristics of the wool itself. CryoSEM gave good preservation of wool and cellular structure but despite numerous random cryo-fractures, an image of a thread exiting the cell through the cell wall gap was never obtained. High resolution light and electron microscopy of embedded and sectioned material gave the best images of the wool exit sites and surrounding subcellular compartments, however, the processing removed the wool threads (although dense vacuolar staining likely attributed to stores of flavones and derivatives can be seen). Observing the gaps within the cuticle, cell wall, plasma membrane and cytoplasm where the thread is presumed to exist, before processing, does give us clues as to how the wool is made; we speculate that a small void in the wall, arising by digestion of the wall constituents, begins on the side of the wall facing the cell (Fig. 5). Flavone material from the vacuole is transported to and deposited into this void. Completion of digestion to include the cuticle permits the flavone aggregate to then extrude outside the cell to form the wool while forming a perfect seal with the edge of the hole (Fig. 5), helped by the large amounts of epicuticular wax. For thread elongation, flavone is continually deposited to the cytoplasmic (elongating) end and the thread can protrude deeper into the cell towards the vacuole(s). This apparent vacuole fusion may arise due to rapid flavone transport and deposition across the tonoplast.

**Figure 5.**
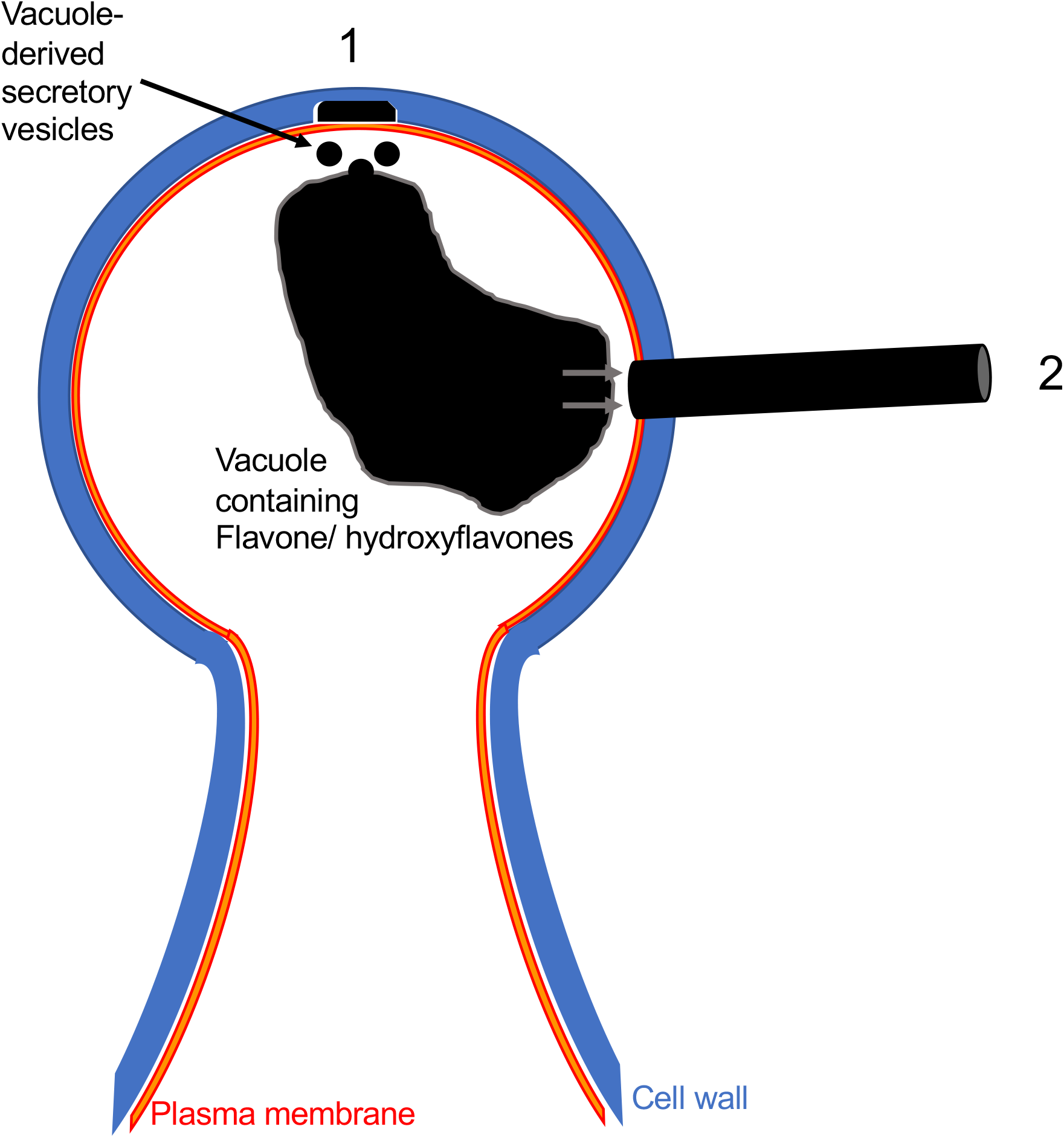
Proposed schematic of farina wool formation by the glandular trichome. (1) Localised cell wall digestion commences from the membrane face and is coupled with deposition of (hydroxy)flavones within the space. (2) Cell wall digestion produces a hole through which the (hydroxy)flavones are extruded. The (hydroxy)flavones may be deposited directly from the vacuole.

A question remains as to how the wooly threads of farina are formed and stably maintained where other flavone-based farina forms a powder. Our compositional analyses suggest the *D. tapetodes* wool is predominantly flavone mixed with 2’- and 4’-hydroxyflavones plus small quantities of other unidentified substituted flavones. The three identified flavones are not unique to this species, since they are found among the powdery *Primula*, where they are referred to as epicuticular flavonoids (Colombo *et al*. 2014). There are, however, data supporting strong intermolecular H-bonding for the 2’ and 4’-hydroxy positions compared to weak H-bonds for the 6-hydroxy position and *intra*molecular bonds in 5-hydroxyflavone (Looker and Hanneman 1962; Looker *et al*. 1966). These intermolecular interactions between the 2’ and 4’-hydroxyflavones and with the bulk flavone may be enough to maintain the integrity of a wool thread. Our Raman data of the *P. marginata* powder show a match with only flavone with no evidence of a mixture like that observed for *D. tapetodes* and so in the former species promoting powder rather than wool extrusion seen in the latter. Despite this observation, chemical analysis of exudate flavonoids demonstrate large quantities of flavone and 2’-hydroxyflavone in *P. marginata* (Valant-Vetschera *et al*. 2009). We put forward a hypothesis that, while a diversity of substituted flavones is maintained across powdery and wooly farina forming species, only wooly farina species can correctly mix and incorporate these at a single site of synthesis to make continuous threads. The selection pressure(s) that have resulted in powdery vs wooly vs efarinose species is not understood, however, the wool does appear to give better coverage over the whole plant while we speculate that the fine loose powder, while more local and easier to dislodge, can yield a denser coating. In addition to the reported capacity of flavonoids to provide protection against UV and potentially during freezing events, the unique capacity of *D. tapetodes* for threading flavones may provide another evolutionary step towards previously described xeromorphic adaptation to drought in the genus *Dionysia* (Wendelbo 1971). Indeed, such specific spatial deposition of flavones seems to offer more leaf coverage from a single production point (glandular trichome), which may limit air movement on the leaf surface, therefore limiting water loss. It remains to be seen whether the properties of wooly farina might make it a useful industrial biomaterial.

## Acknowledgements

We thank Trevor Groves with help with low kV SEM imaging of uncoated fibres and Prof. Beverley Glover for critical reading of the manuscript. We are grateful for the opportunity to access the *Dionysia tapetodes* and *Primula marginata* accessions at Cambridge University Botanic Garden. TEM and FE-SEM were performed using the EM facilities of the Cambridge Advanced Imaging Centre (CAIC), University of Cambridge. The Spring lab acknowledges support from EPSRC, BBSRC and the Royal Society. JG thanks the BBSRC Doctoral Training Programme and AstraZeneca for funding. The Microscopy Core facility at the Sainsbury Laboratory is supported by the Gatsby Charitable Foundation.

## Supplementary Data list

**Dataset S1 comprising Figures S1-S7 and Table S1.** All data and interpretations/ commentary for HLPC, LCMS, HMRS and NMR chemical analyses.

**Figure S8**.

**Figure S9.**

## Dataset S1 including Figures S1 to S7 and Table S1

### Analytical HPLC analysis of wool fibre samples

To assess the purity of the sample, HPLC analysis was performed using two different gradients (5-95% and 40-95% acetonitrile in water) to ensure that all peaks present were observed.

**Figure S1a.**
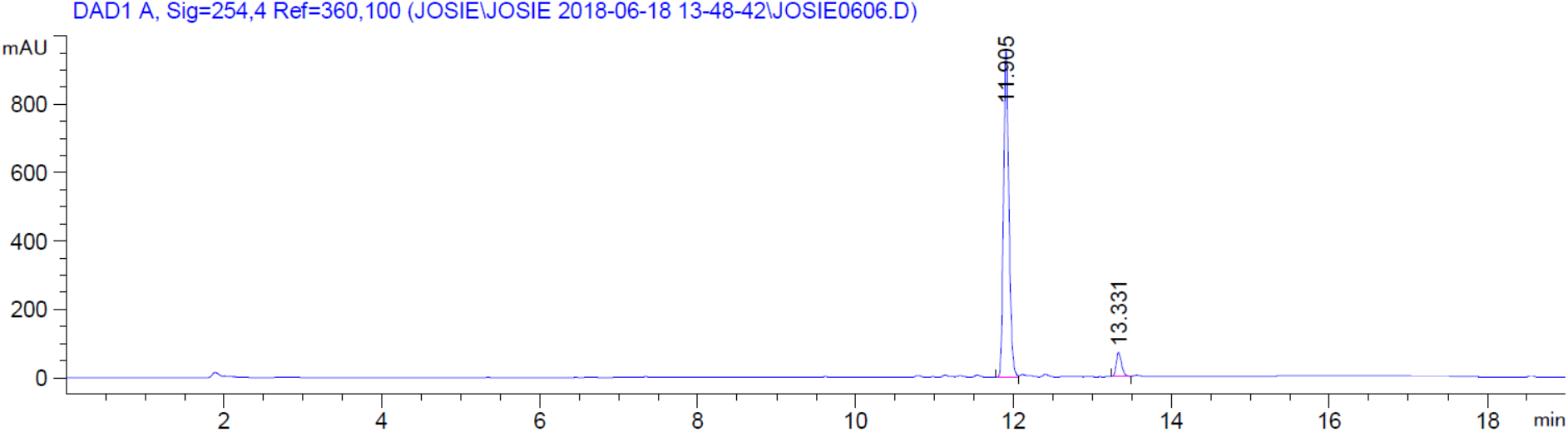
The HPLC spectra for analysis of the sample using a 5-95% acetonitrile in water gradient. The large signal at 2-2.5 minutes in the 220 nm trace (Figure 1 and 2) is due to dimethylsulfoxide (DMSO) which was used to dissolve the sample.

**Figure S1b.**
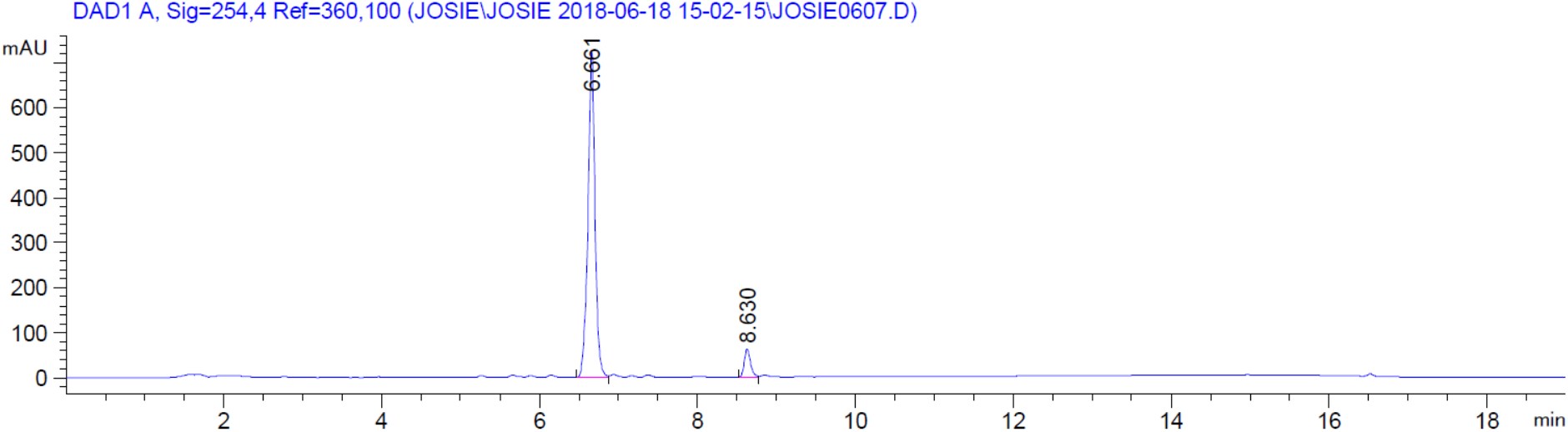
The HPLC spectra for analysis of the sample using a 5-95% acetonitrile in water gradient.

**Table S1.** The peaks observed in the spectra shown in Figure 1.

| Peak | Retention time (min) | Area (mAU*s) | Area (%) |
| --- | --- | --- | --- |
| 1 | 11.905 | 4507.078 | 93.0 |
| 2 | 13.331 | 340.296 | 7.0 |
| 1 | 6.661 | 4535.346 | 93.0 |
| 2 | 8.630 | 339.714 | 7.0 |

### S2: Mass spectrometry of wool fibre sample

**Figure S2a.**
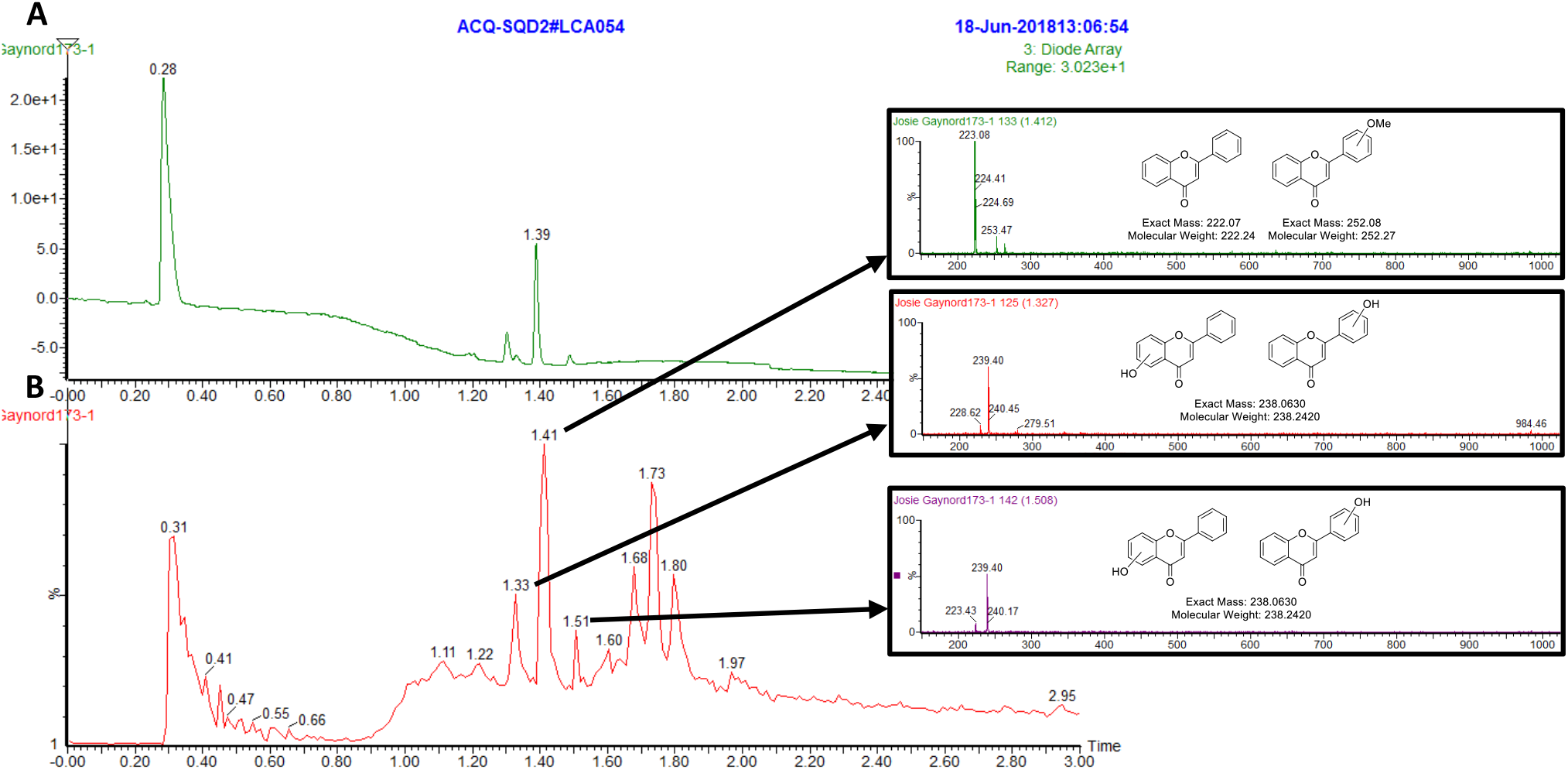
A: The UV trace for the LCMS spectrum of the plant sample. B: The corresponding ES+ trace. Insets are mass traces (ES+) for individual peaks, as indicated by the arrows. Suggested chemical structures with exact masses and molecular weights are shown over the relevant mass spectra.

**Figure S3.**
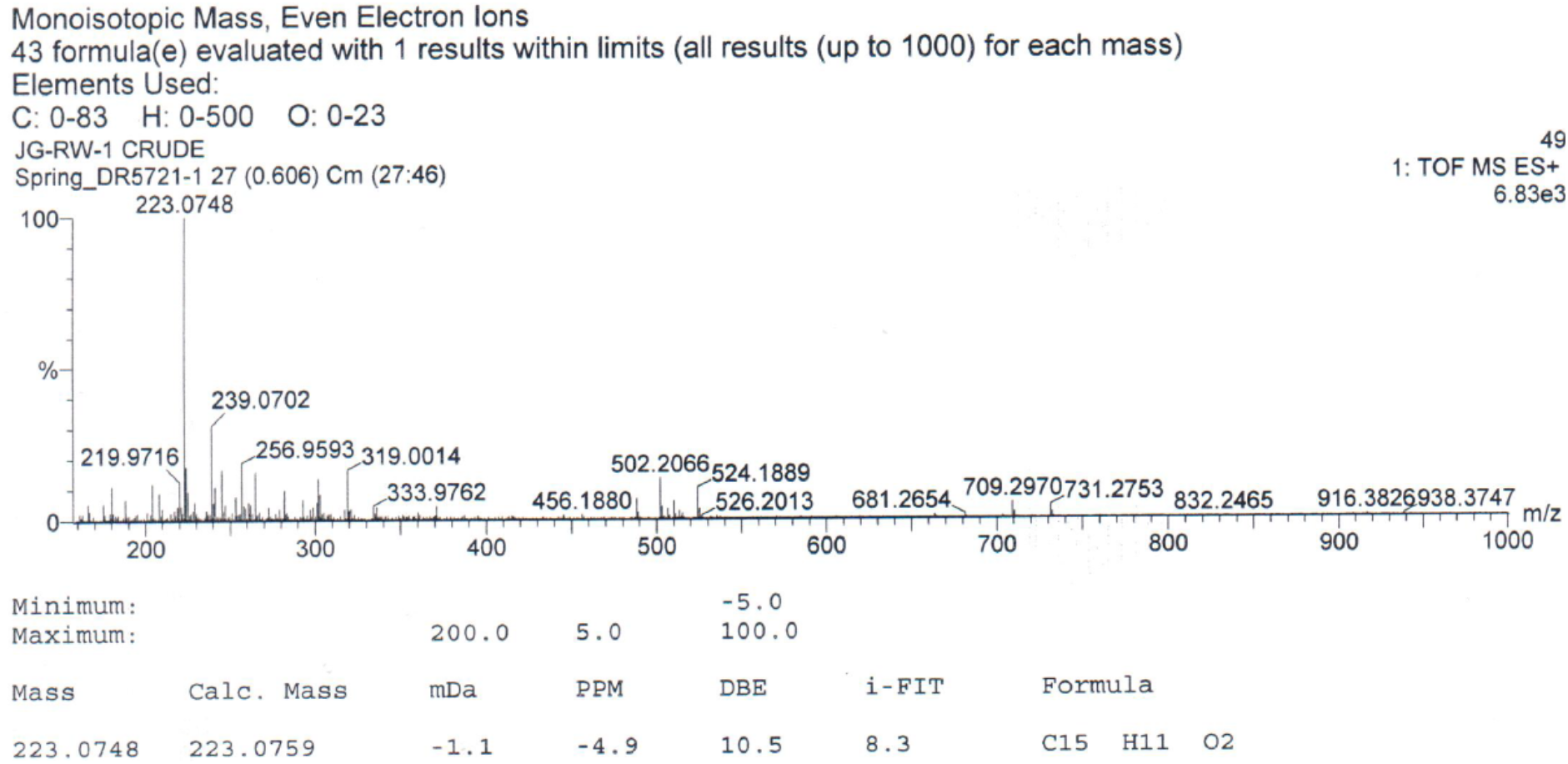
The high-resolution mass spectrum (HRMS) for the plant sample.

### HPLC analysis of pure flavone

**Figure S4.**
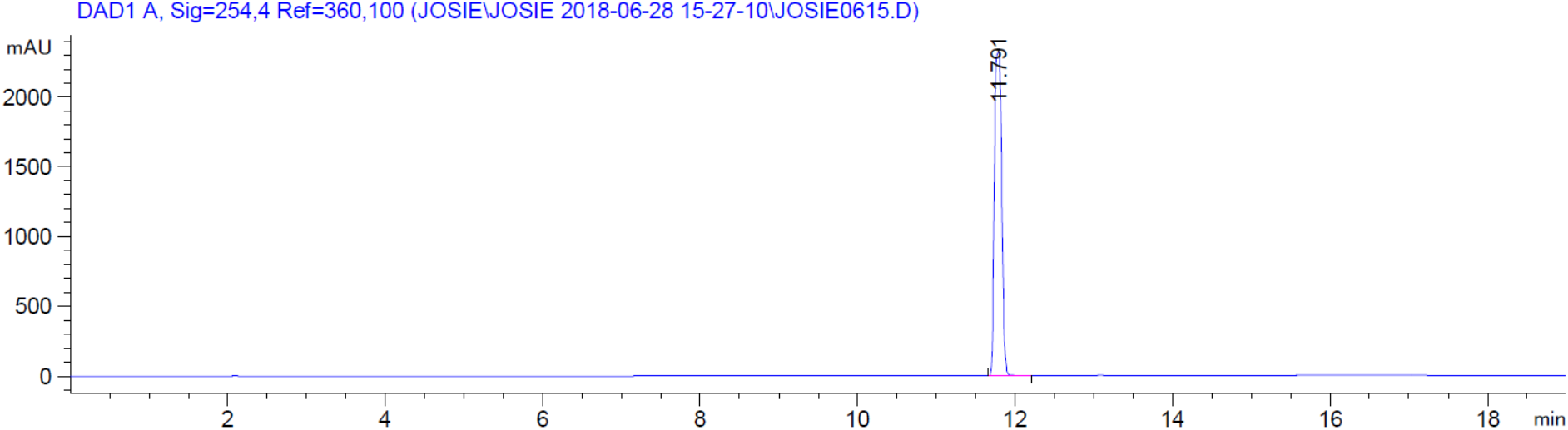
The HPLC spectra for analysis of pure Flavone using a 5-95% acetonitrile in water gradient.

### NMR comparison of commercially-available flavone and wool fibre

**Figure S5a.**
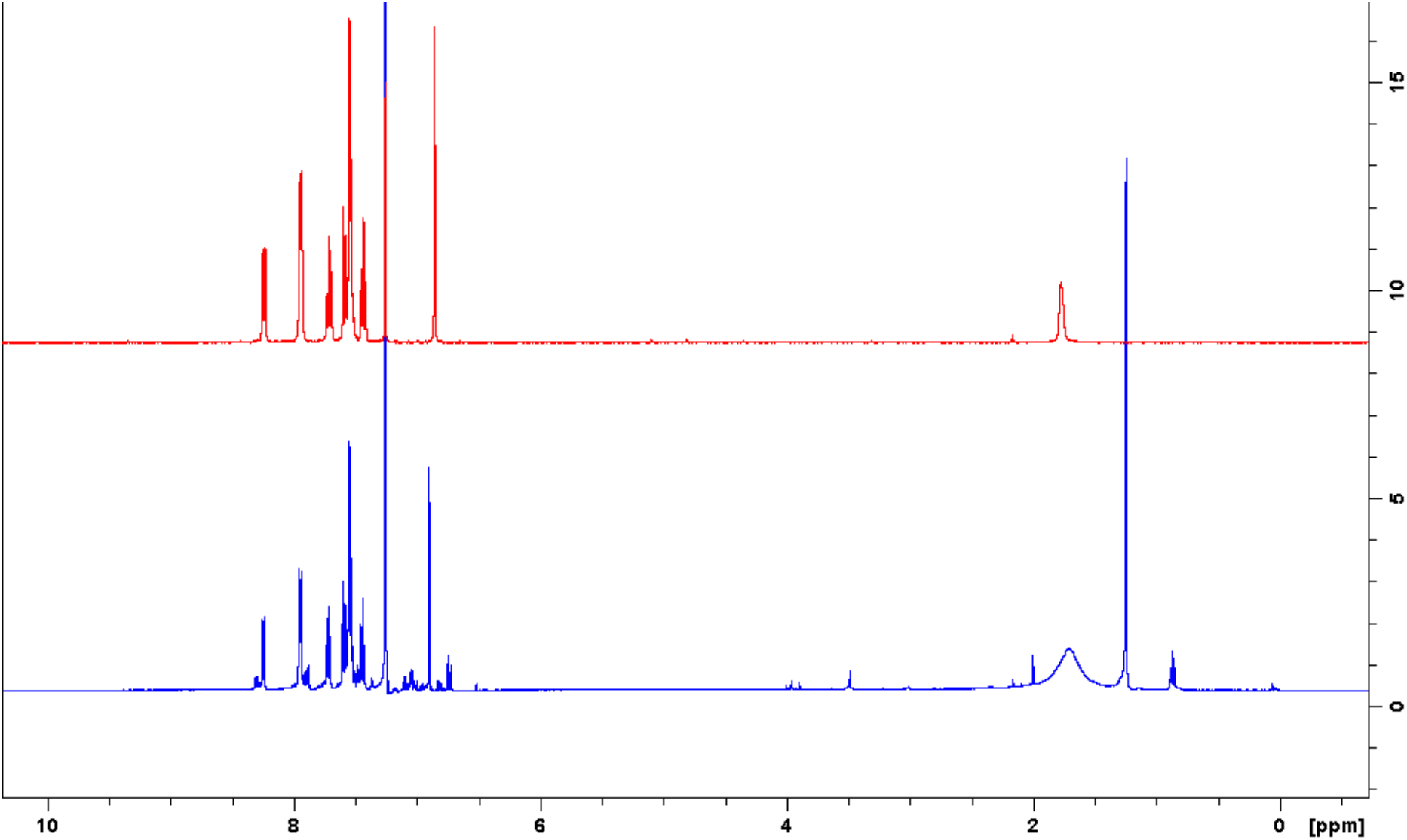
A comparison of the ^1^H NMR spectra for a pure sample of flavone (red) and the wool fibre sample (blue) between 10 and 0 ppm.

**Figure S5b.**
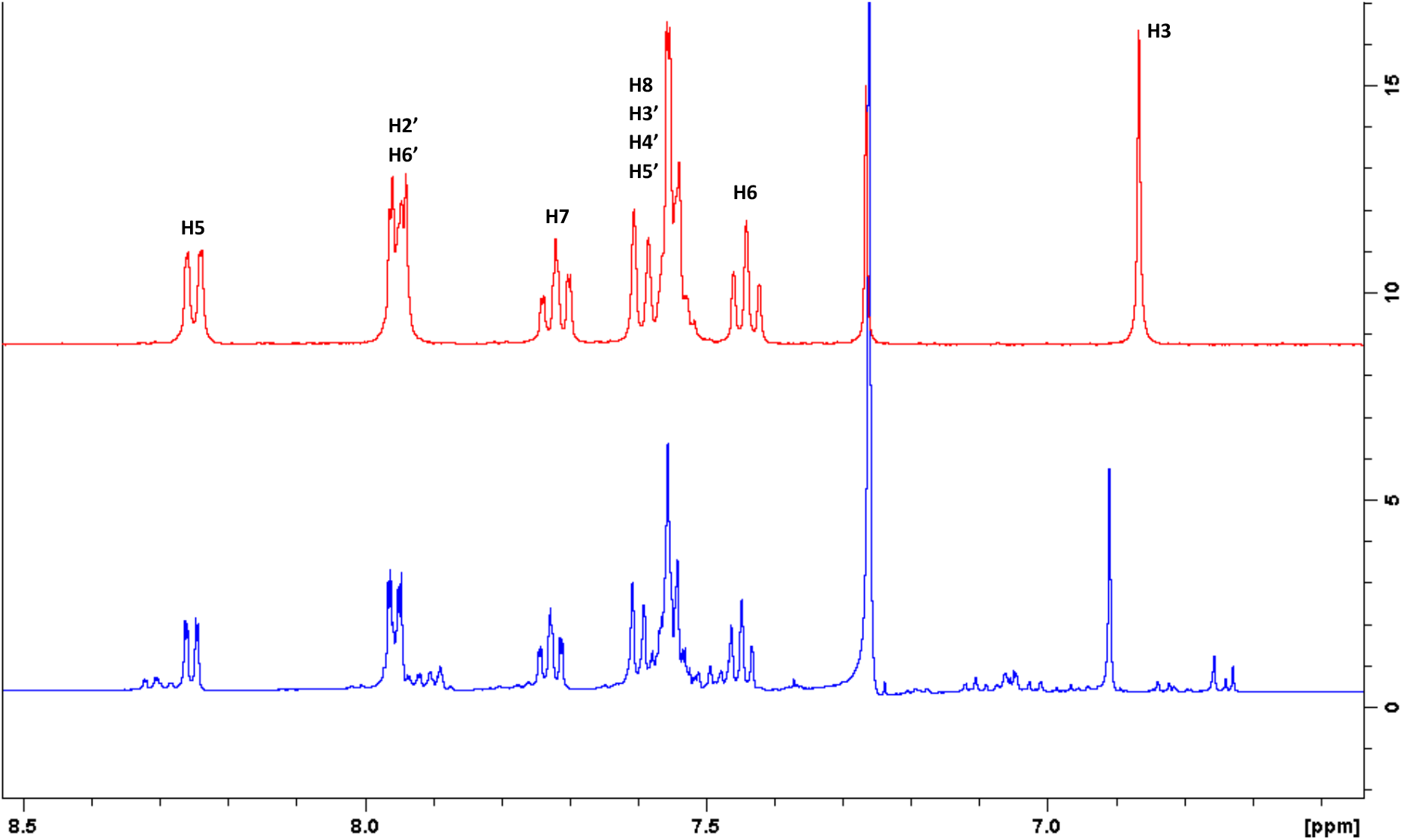
A comparison of the ^1^H NMR spectra for a pure sample of flavone (red) and the wool fibre sample (blue) between 8.5 and 6.5 ppm.

**Figure S5c.**
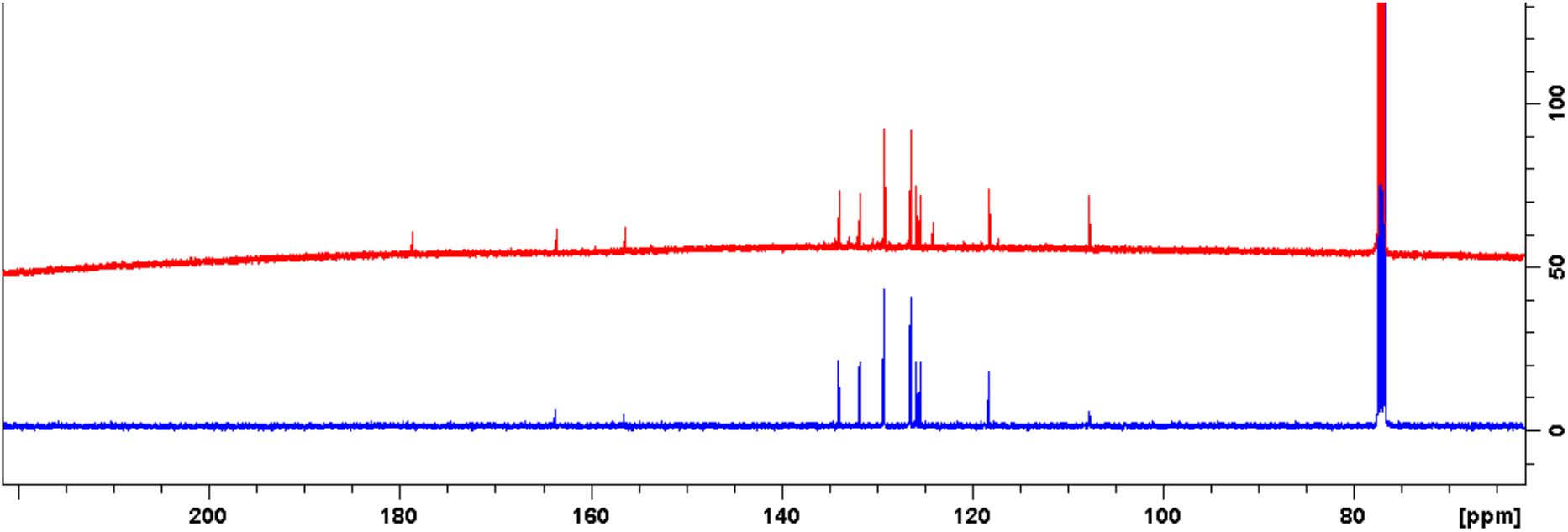
A comparison of the ^13^C NMR between 220 and 70 ppm for a pure sample of flavone (red) and the wool fibre sample (blue).

### NMR Analysis of wool fibre sample

The sample was analysed using the following NMR experiments: ^1^H NMR (Figure S6a), ^13^C NMR (Figure S6b), DEPT 135 (Figure S6c), COSY (Figure S6d), HSQC (Figure S6e) and HMBC (Figure S6f).

The ^1^H and ^13^C NMR of pure flavone were compared to the plant sample and the signals were found to match (S5), providing further evidence that the major component of the plant sample mixture is unsubstituted flavone. The NMR assignments are therefore as follows^1,2^:

**^1^H NMR** (500 MHz, CDCl3) δ 8.26-8.24 (m, 1H, H5), 7.96-7.94 (m, 2H, H2’, H6’), 7.74-7.71 (m, 1H, H7), 7.60-7.51 (m, 4H, H8, H3’, H4’, H5’), 7.46-7.43 (m, 1H, H6), 6.91 (s, 1H, H3)
**^13^C NMR** (125 MHz, CDCl3) δ 178.7 (C, C4), 163.6 (C, C2), 156.5 (C, C9), 134.0 (CH, C7), 131.9 (C, C1’), 131.8 (CH, C3’, C5’), 129.2 (CH, C4’), 126.5 (CH, C2’, C6’), 125.9 (CH, C5), 125.4 (CH, C6), 124.1 (C, C10), 118.3 (CH, C8), 107.8 (CH, C3)

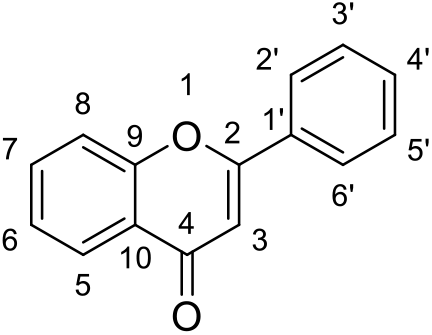

**Figure S6a.**
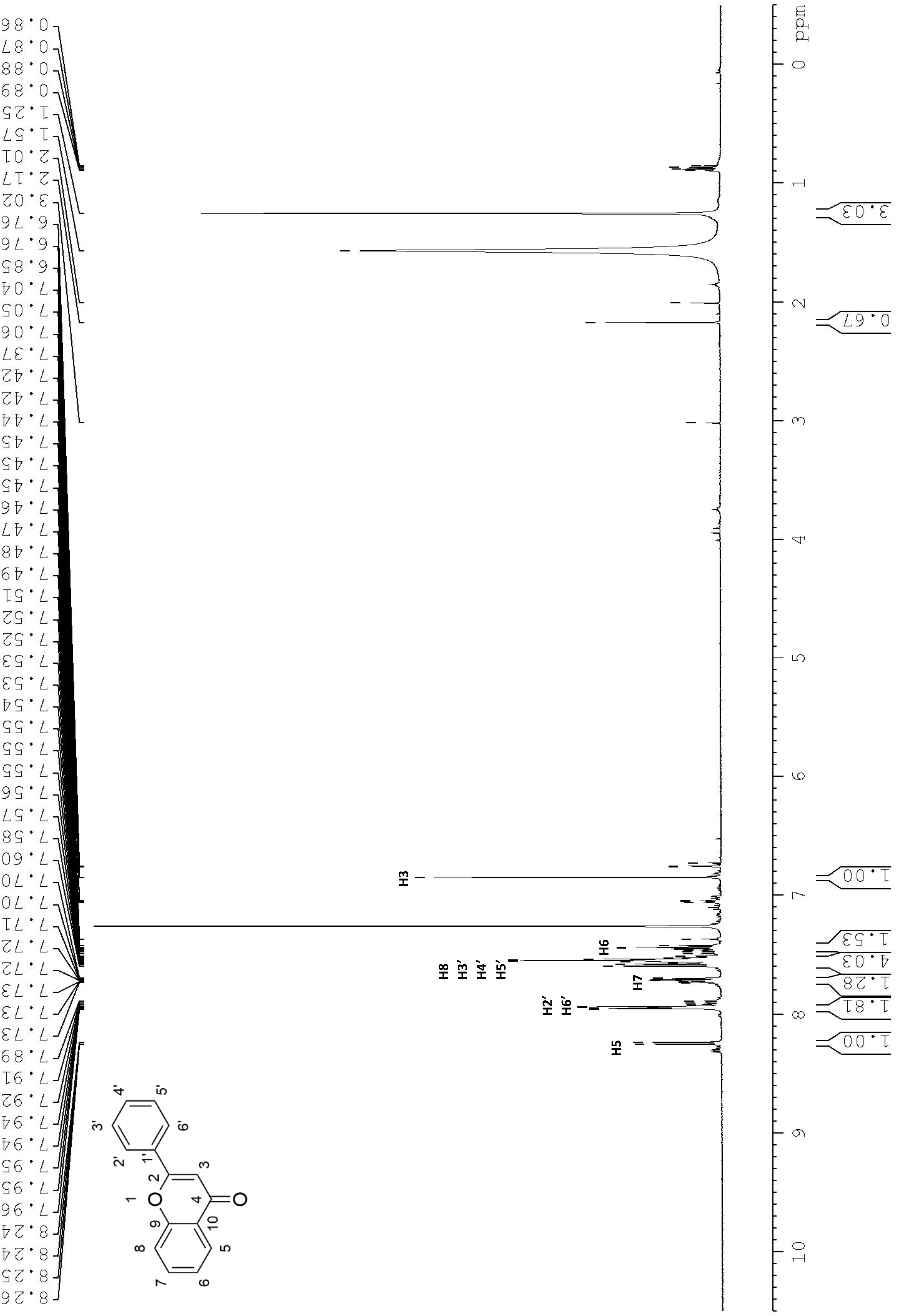
The assigned ^1^H NMR spectrum of the plant sample.

**Figure S6b.**
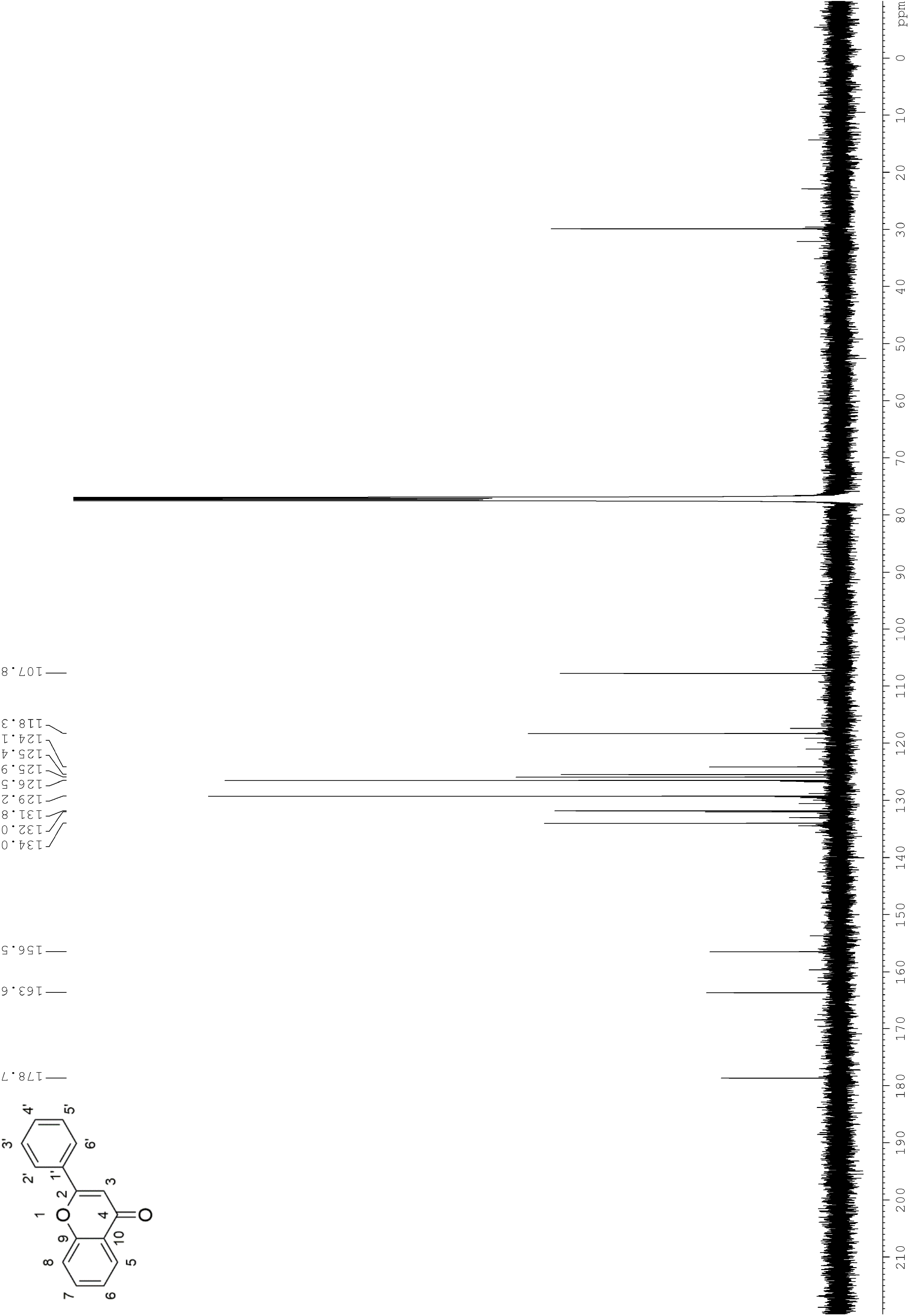
The ^13^C NMR spectrum of the plant sample

**Figure S6c.**
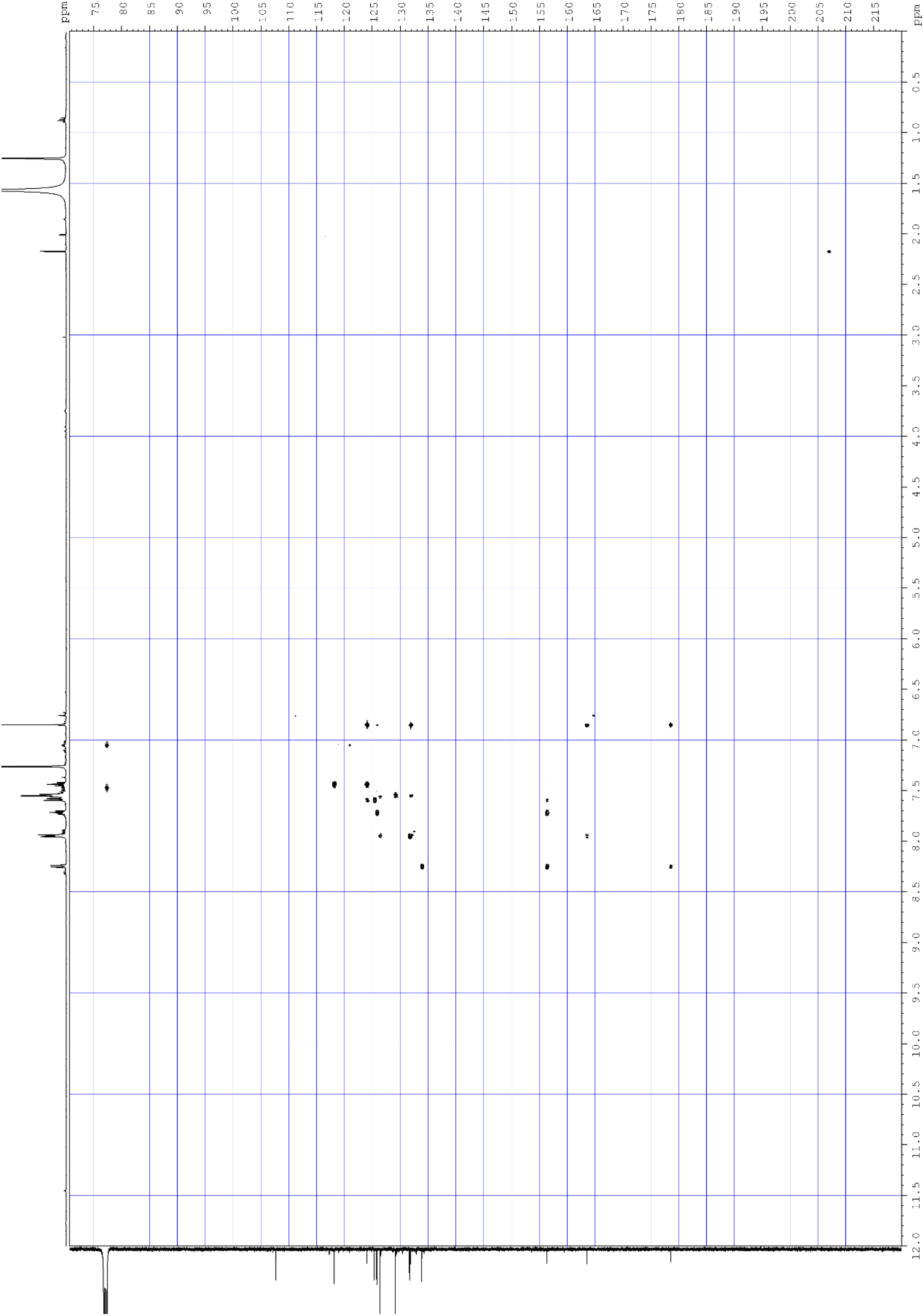
The HMBC spectrum of the plant sample

**Figure S6d.**
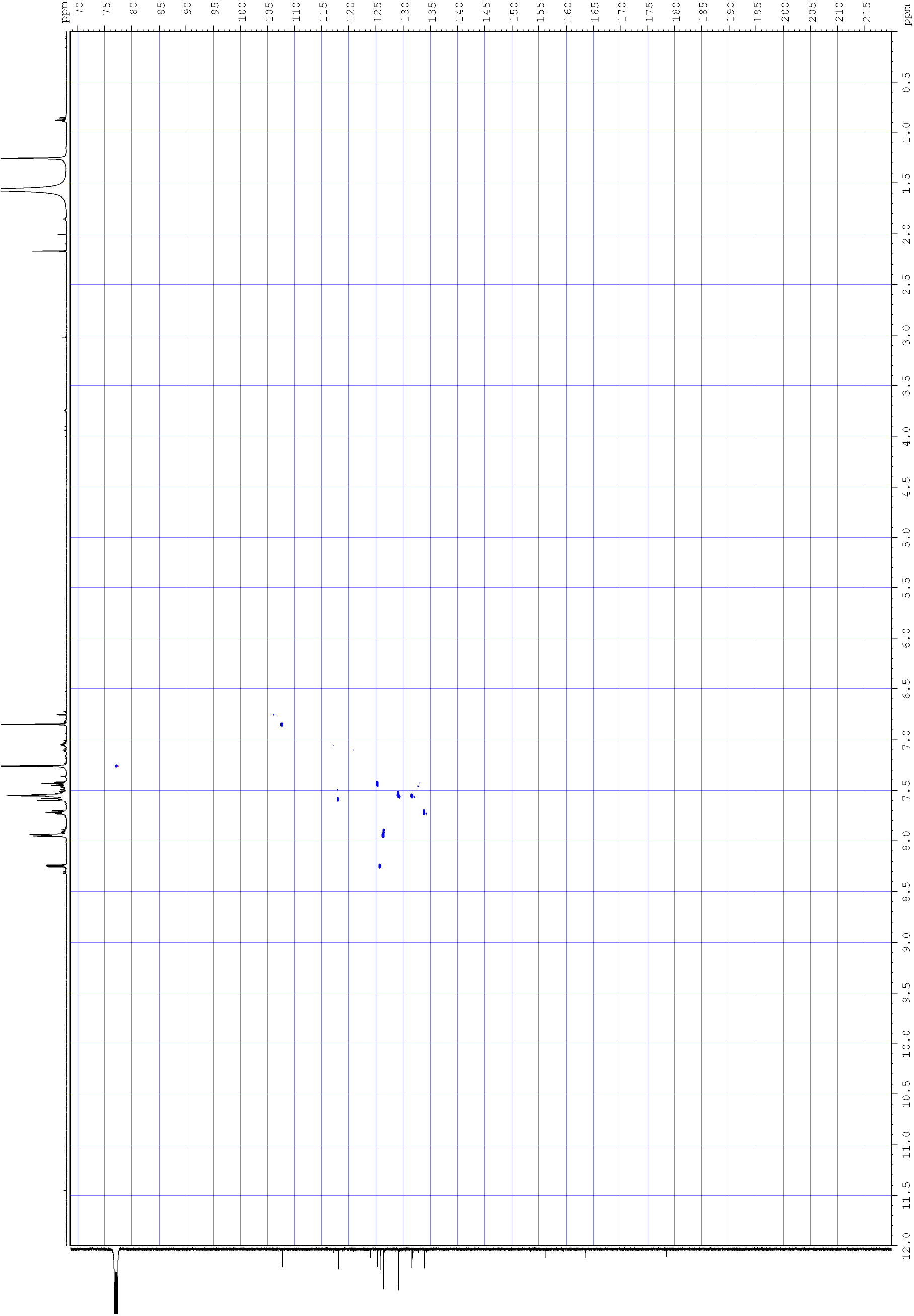
The HSQC spectrum of the plant sample

**Figure S6e.**
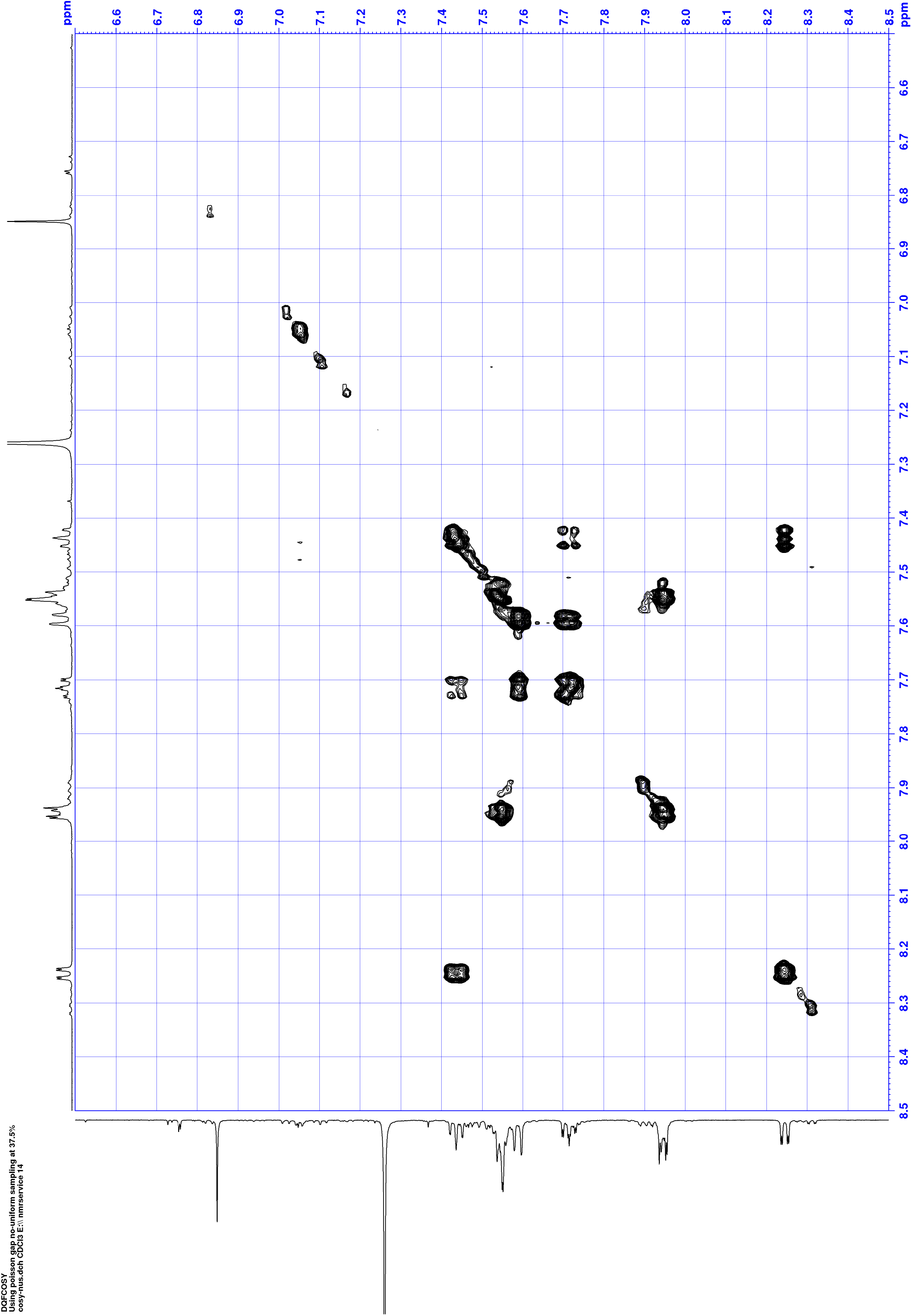
The COSY spectrum of the plant sample

**Figure S6f.**
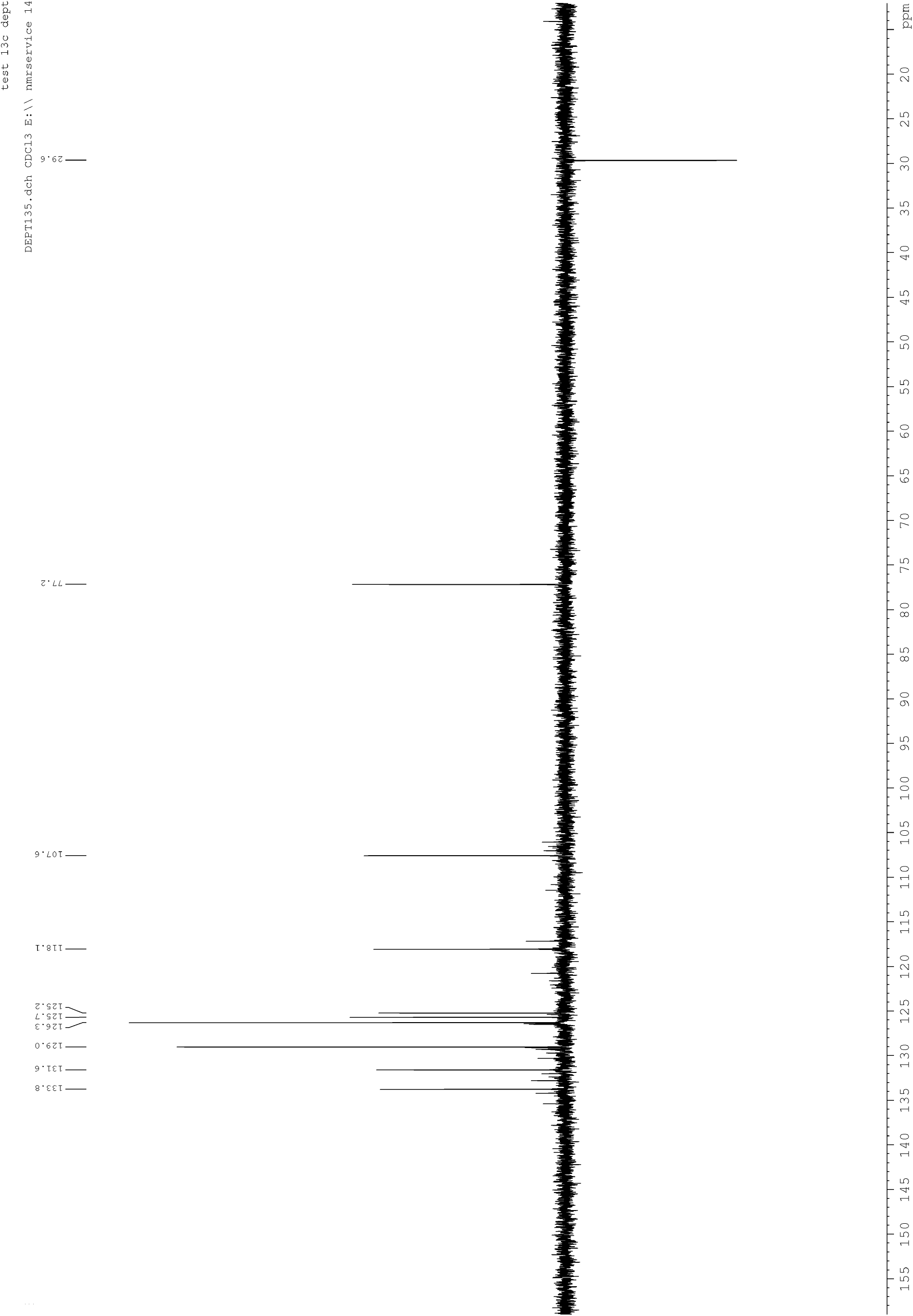
The DEPT 135 spectrum of the plant sample

### 2D NMR experiments to investigate substitution patterns on flavones

To investigate the structure of the minor species present in the wooly farina sample, a HSQC experiment was run with non-uniform sampling to increase the resolution of the spectrum (Figures S7a and S7b).

**Figure S7a.**
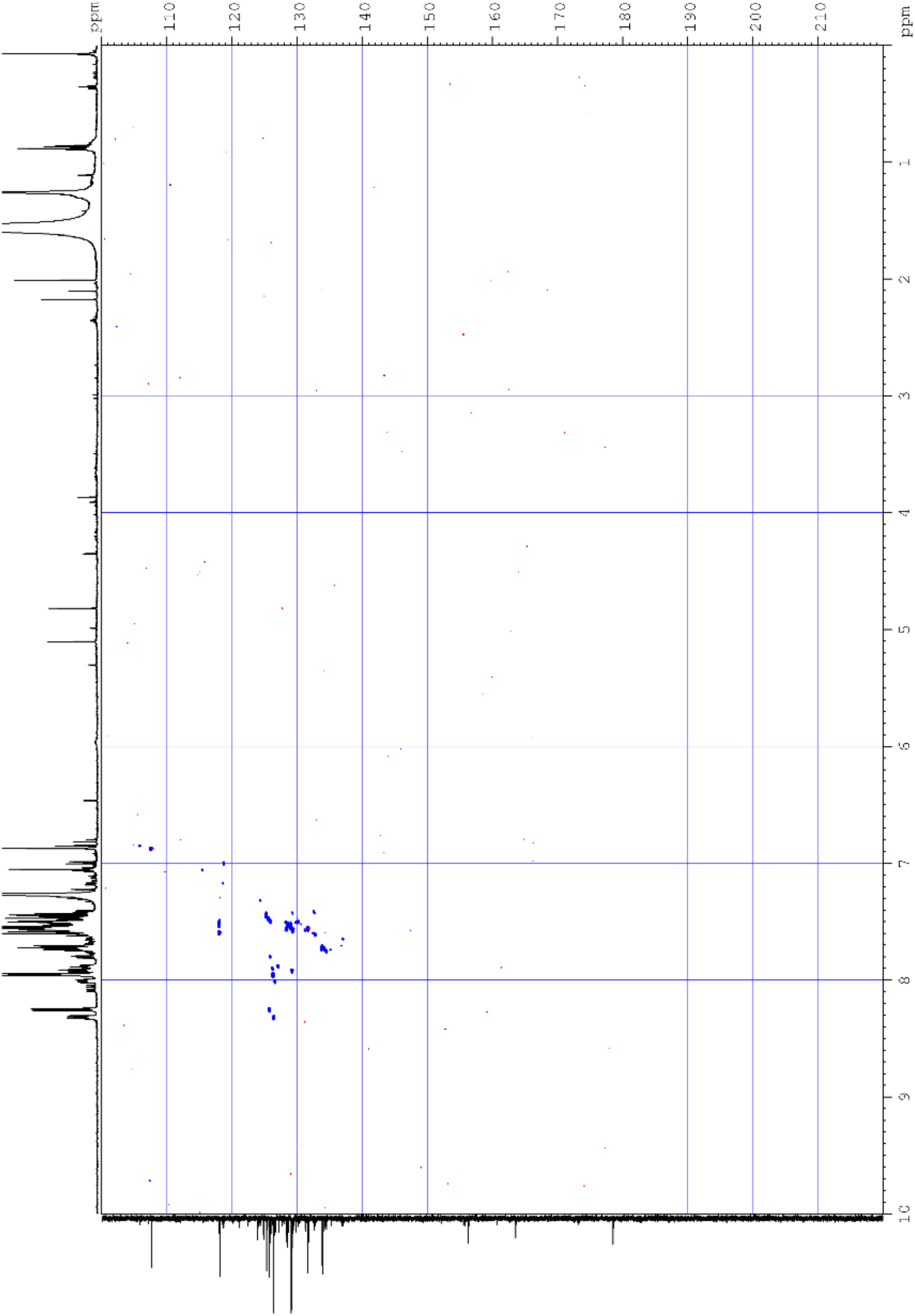
The spectrum of the HSQC experiment (run with non-uniform sampling).

**Figure S7b.**
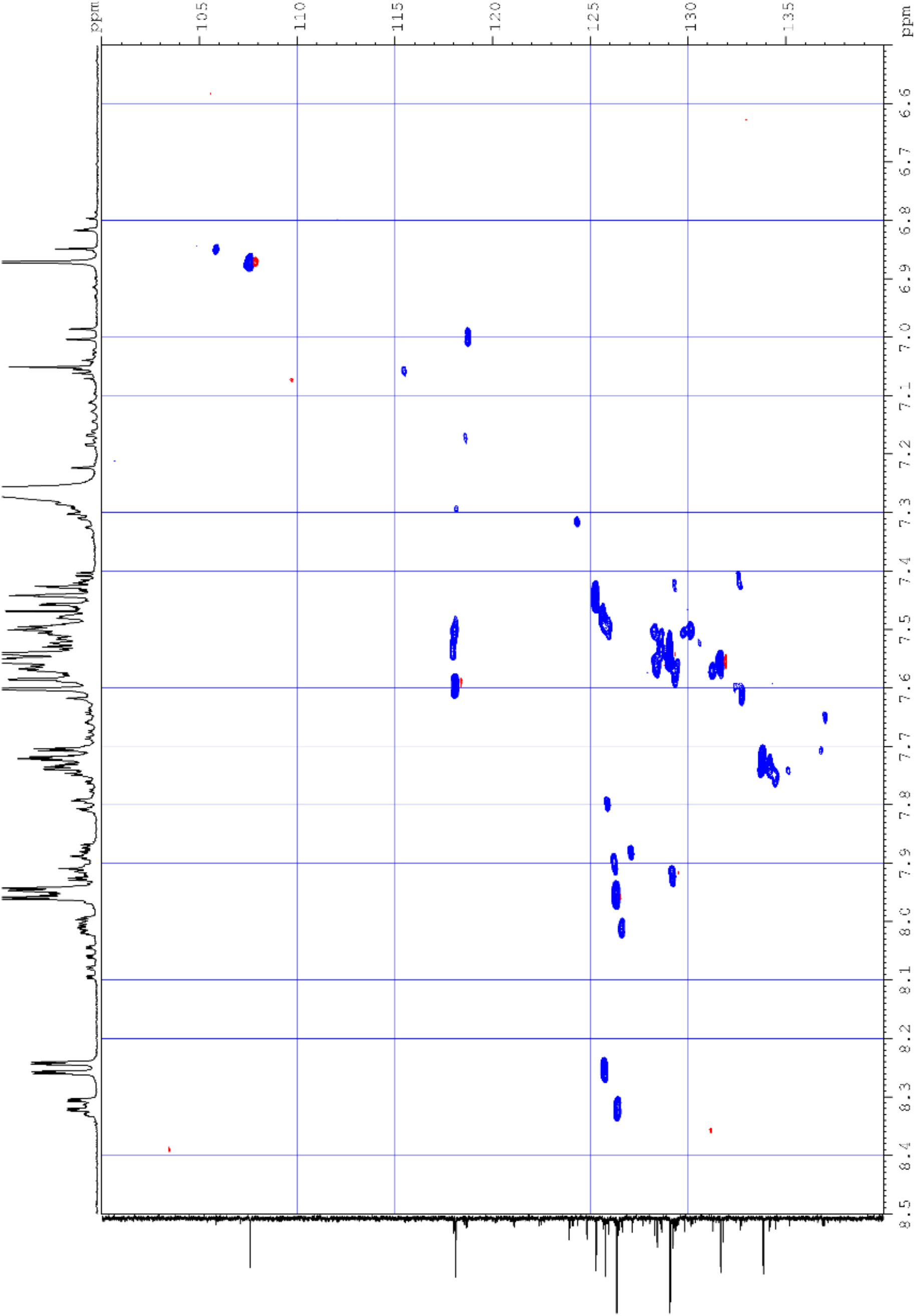
A zoomed in section of the aromatic region of the HSQC experiment (run with non-uniform sampling).

Without NMR, LCMS, HRMS and HPLC analysis of the isolated components of the plant sample mixture, the structure of the minor species cannot be stated with 100% confidence. However with the data from the LCMS, HPLC, HRMS and NMR, suggestions can be made.

The masses from the LCMS indicated that mono-hydroxy or mono-methoxyflavones were present in the wooly farina sample. Literature values for carbon atoms bearing a hydroxy or methoxy substituent in flavones are very similar and signals are present at 3.9-4.0 ppm in the ^1^H NMR spectrum which could correlate to the methyl group of a methoxy substituent, however with the NMR data available it could not be confirmed if methoxy-substituted flavones were present.

For hydroxy substitution, each potential substitution site was considered in turn. The expected shifts of the protons and carbons with hydroxy-substitution were based on literature values^3,4^. From the information provided by the HSQC experiment, the most likely substitution pattern is 4’-hydroxyflavone:

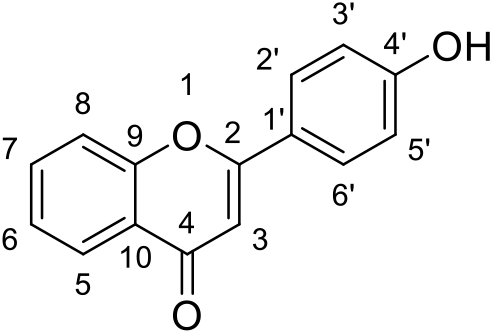

The evidence for this is as follows:

A signal is present in the ^13^C NMR spectrum at 160.7 ppm which is in the correct region for the quaternary C4’ carbon with either a hydroxyl substitution (Figure S7c).

Correlation is observed in the HSQC between a ^13^C signal at 127.1 ppm and a ^1^H signal at 7.88 ppm (A, Figure S7d). These match the expected shifts for an aromatic CH in a meta position to the hydroxylation site (i.e. the CH at either C2’ or C6’).

Correlation is observed between a ^13^C signal at 118.6 ppm and a ^1^H signal at 7.16 ppm (B, Figure S7d) which match the expected shifts for an aromatic CH in an ortho position relative to the hydroxyl group (i.e. the CH at C3’ or C5’).

**Figure S7c.**
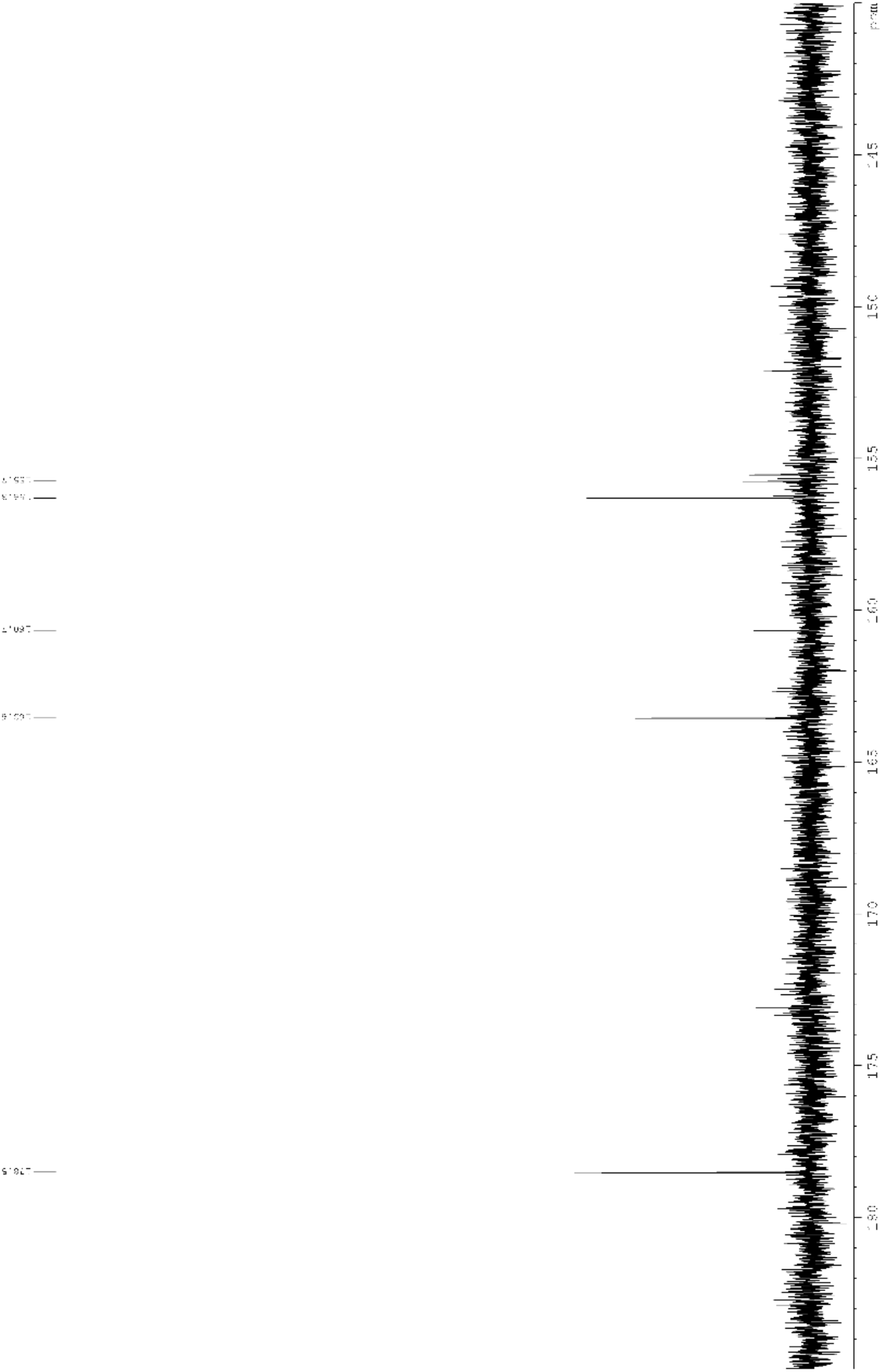
The ^13^C NMR spectrum showing the peak at 160.3 ppm

**Figure S7d.**
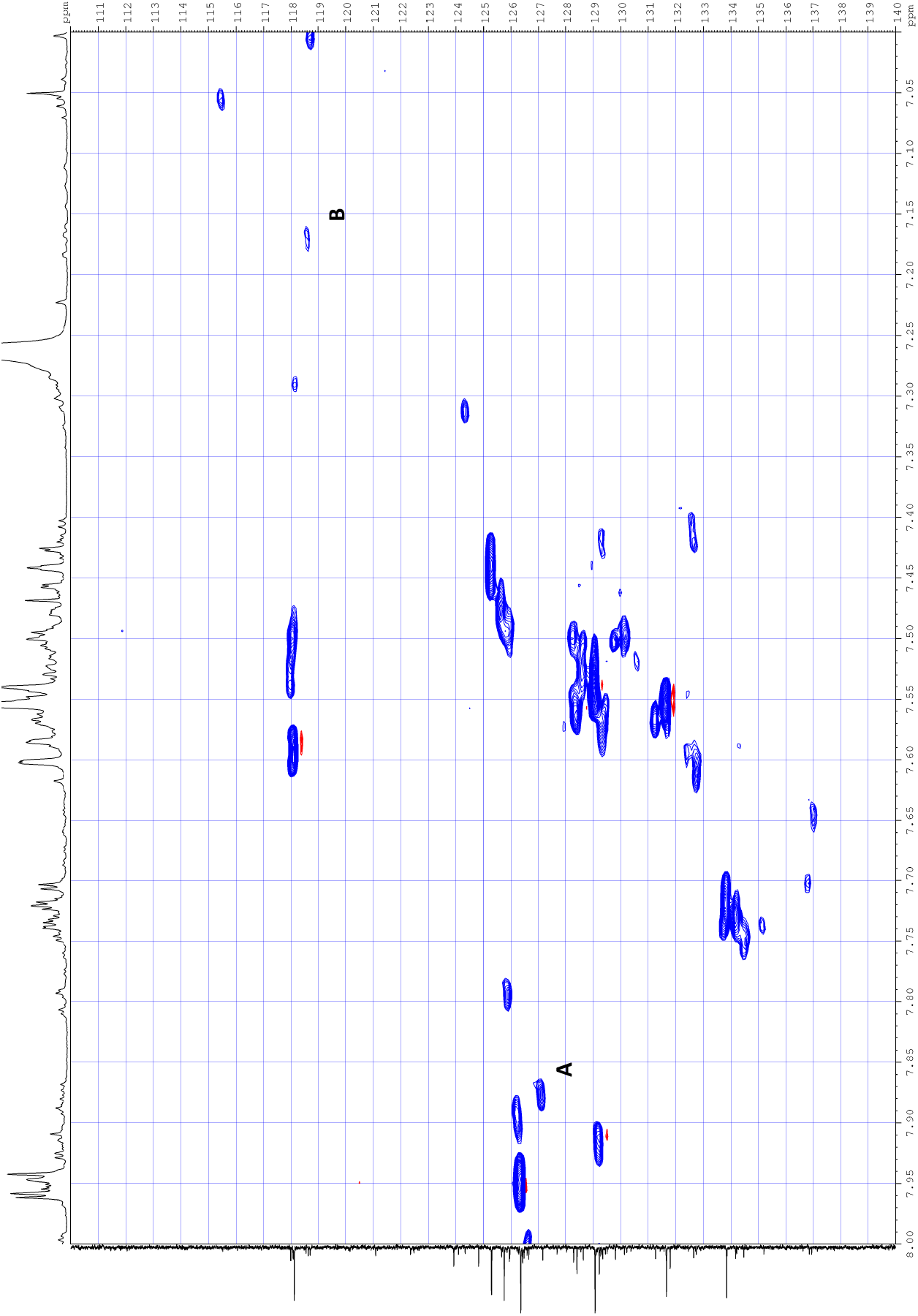
A zoomed in region of the HSQC (with non-uniform sampling) showing the two correlations A and B.

As the LCMS data indicated there were two mono-hydroxylated species with different retention times present in the plant sample, there could be another isomer present. The NMR data point towards this being 2’-hydroxyflavone:

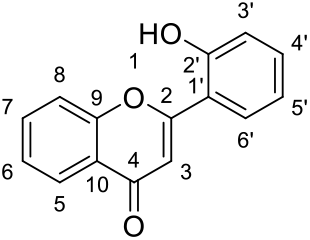

The evidence for this includes:

In the ^13^C NMR spectrum there is a peak at 152.1 ppm, which is around the expected shift for the quaternary C2’ bearing the hydroxyl group (Figure S7e).

In the HSQC experiment with non-uniform sampling, a peak at 132.6 ppm in the ^13^C NMR correlates with a peak at 7.41 ppm in the ^1^H NMR (A, Figure S7f). These shifts match the expected signal for an aromatic CH which is in a meta position relative to the hydroxyl (i.e. the signal for H4’)

In the HSQC experiment with non-uniform sampling, a peak at 124.3 ppm in the ^13^C NMR correlates with a peak at 7.31 ppm in the ^1^H NMR (B, Figure S7f). These shifts match the expected signal for an aromatic CH which is in a para position to the hydroxyl group (i.e. for C5’).

**Figure S7e.**
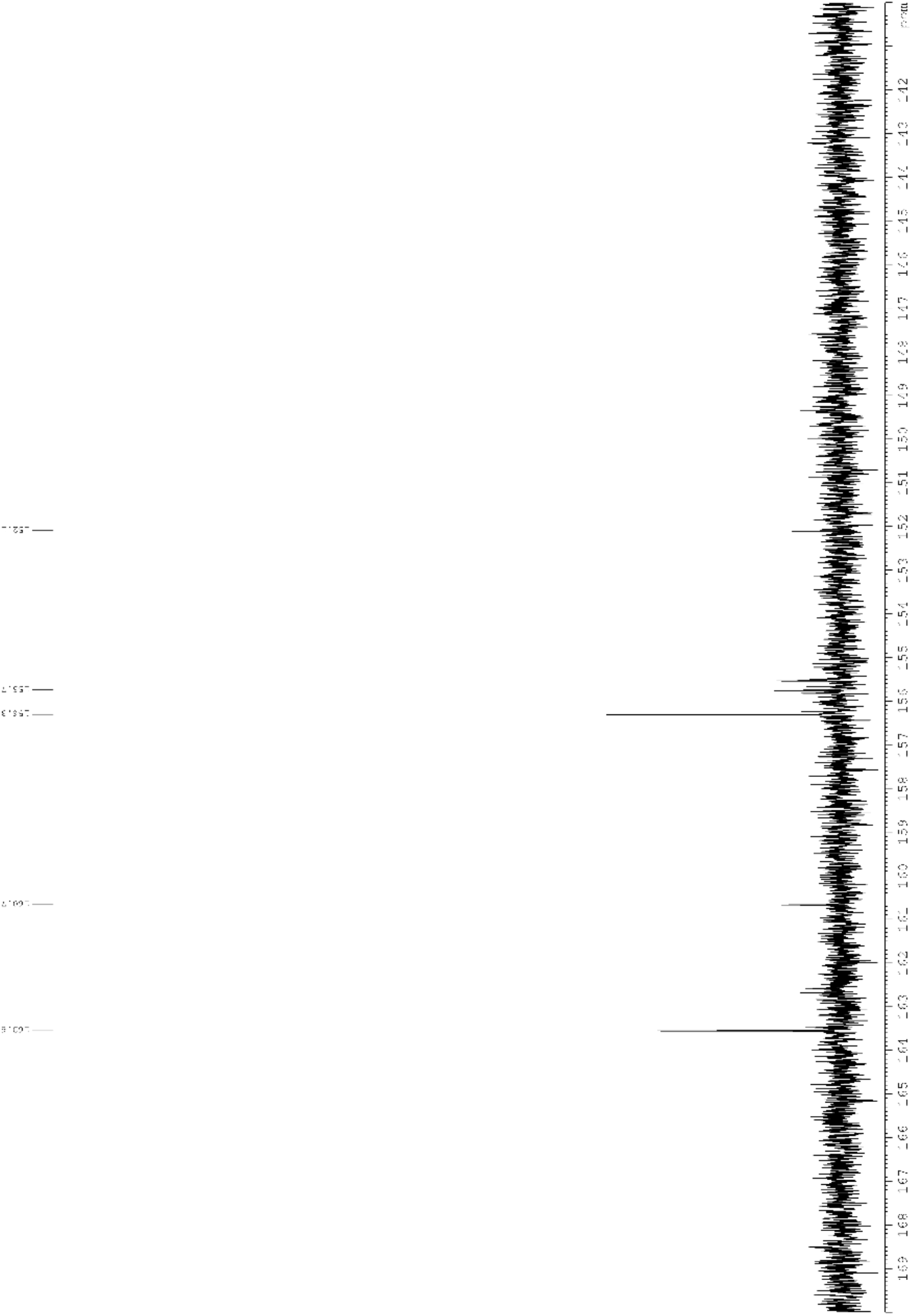
A zoomed in region of the HSQC (with non-uniform sampling) showing the two correlations A and B.

**Figure S7f.**
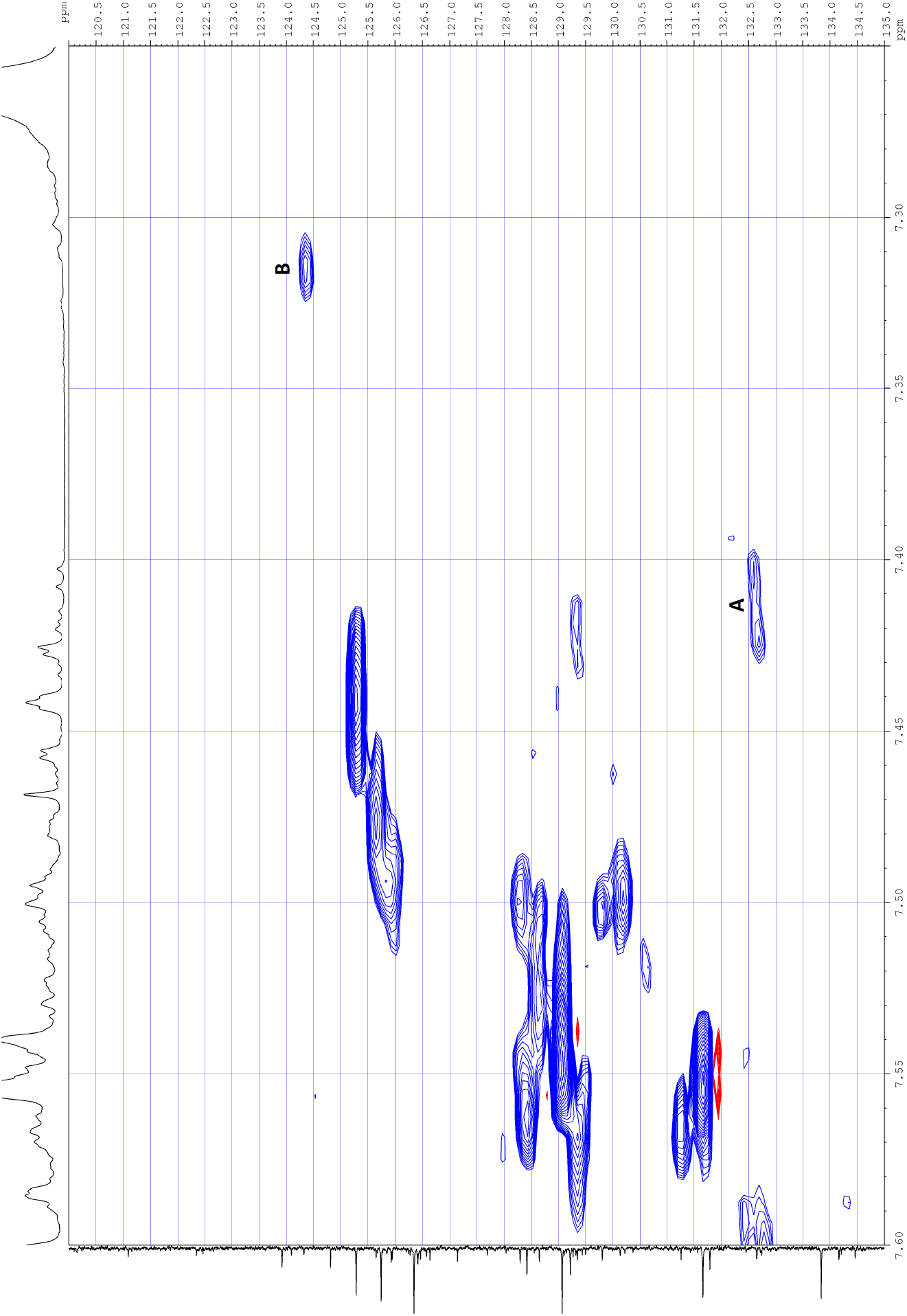
A zoomed in region of the HSQC (with non-uniform sampling) showing the two correlations A and B.

**Figure S8.**
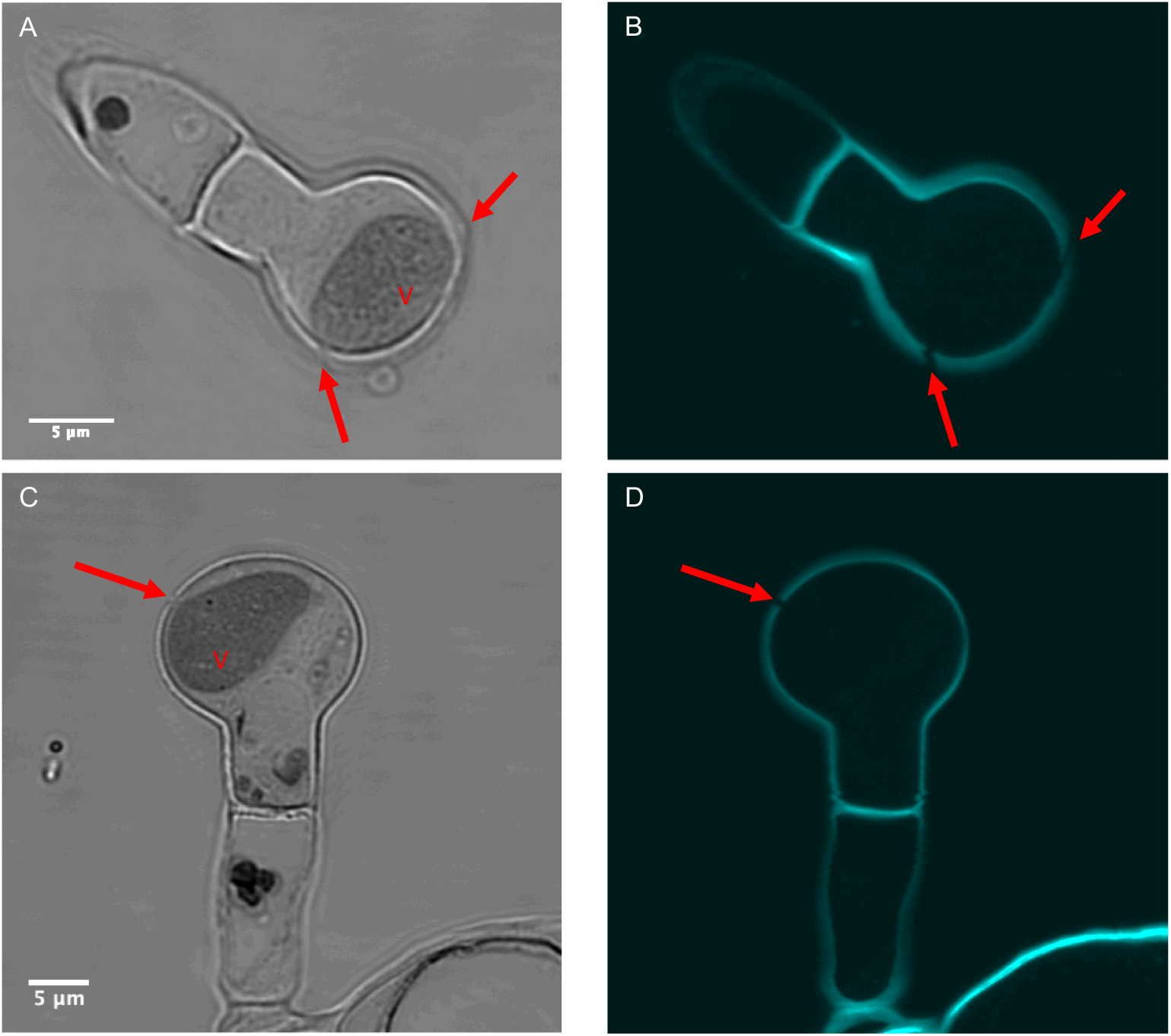
Confocal transmitted image (A, C) and cell wall fluorescence (B, D) of calcofluor-stained sections through gland hair cells. Wool exit holes, observed as discrete gaps in the fluorescence images (arrows) are in close proximity to the dense vacuole (V).

**Figure S9.**
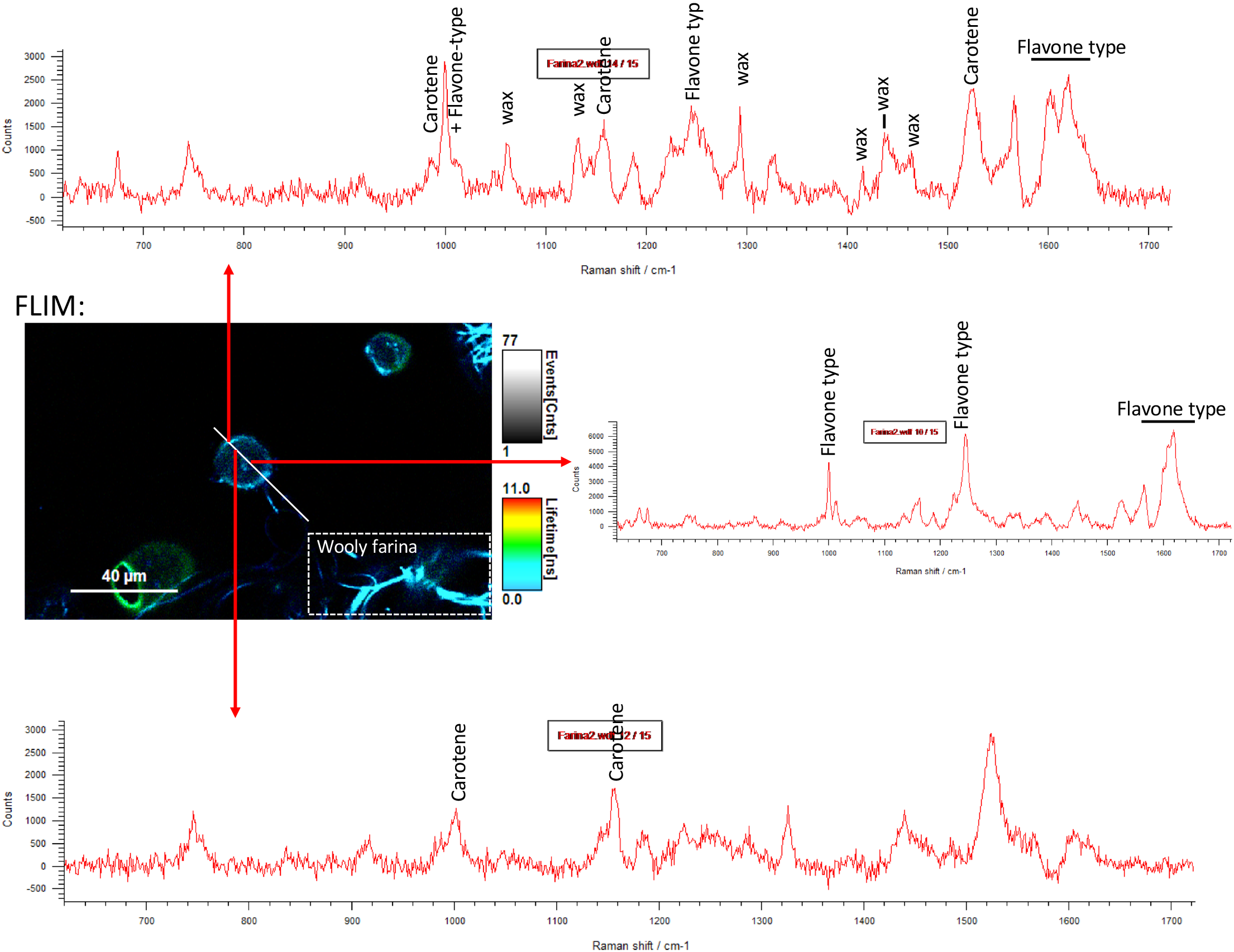
Fluorescence Lifetime Imaging (FLIM) of glandular trichomes taken together with Raman spectra acquired along a line through a trichome cell. The FLIM data (centre left image) represents lifetimes of autofluorescence. Farinose material including the wool and edge of the trichome cell have short lifetimes (cyan and blue colours) with high signal and the cell interior has blue (short) and green (long) lifetimes with low signal. Raman measurements at precise locations along the line confirm the presence of flavone-type material at the cell edge together with strong peaks equivalent to those of plant epicuticular wax (Upper spectrum, assignments are given for prominent peaks). Note the intense cyan labelling in the FLIM image that is a similar lifetime to the wooly farina (white boxed region). Within a proximal location inside the cell there is an absence of the the wax-associated peaks and the strong flavone peaks (lower spectrum). Carotenoid is detected at both locations. Another location inside the cell yielded strong flavone peaks (centre spectrum) and may represent an intracellular store of flavones.

